# Experience-dependent plasticity of a highly specific olfactory circuit in *Drosophila melanogaster*

**DOI:** 10.1101/2023.07.26.550642

**Authors:** Benjamin Fabian, Veit Grabe, Rolf G. Beutel, Bill S. Hansson, Silke Sachse

## Abstract

*Drosophila melanogaster* encounters a variety of odor cues signaling potentially harmful threats throughout its life, which are detected by highly specialized olfactory circuits enabling the animal to avoid them. We studied whether such crucial neuronal pathways are hard-wired or can be modulated by experience. Using long-term exposure to high concentrations of geosmin, an indicator of potentially lethal microorganisms, we demonstrate at the single-cell level that the underlying neuronal circuitry undergoes structural changes in the antennal lobe, while higher brain centers remain unaffected. In particular, second-order neurons show neurite extensions and synaptic remodeling after the exposure period, whereas olfactory sensory neurons and glia cells remain unaffected. Flies that were exposed to geosmin tolerate this innately aversive odorant in general choice and oviposition assays. We show that even a highly specific olfactory circuit is plastic and adaptable to environmental changes.

**HIGHLIGHTS:**

- The highly specific geosmin circuit is modulated by experience-dependent plasticity
- PN dendritic extensions cause a volumetric increase of the geosmin-detecting glomerulus
- LNs are remodeled while OSNs and glia cells remain unaffected by long-term exposure
- Flies adapt their behavior to an odorant signaling a potential lethal threat

## INTRODUCTION

It seems widely accepted that the mammalian central nervous system is at least partly able to undergo morphological plastic changes. However, for decades the insect brain was considered as hard-wired (Heisenberg et al., 1995; Jefferis & Hummel, 2006) and plasticity with regard to the insect nervous system was discussed controversially. As one of the experimentally most accessible parts of any insect brain, the olfactory system of *Drosophila melanogaster* Meigen, 1830 is the source of most of our understanding about stereotypy and plasticity in the insect nervous system. Studies that focused on this model system could show that there is evidence for stereotypy in the axonal innervation pattern of projection neurons (PNs) in the lateral horn (LH), a higher brain center involved in mediating innate behavior (Schultzhaus et al., 2017). Comparisons of PN axon terminals could demonstrate that PNs that innervate the same glomerulus show very similar axonal branching patterns (Jefferis et al., 2007; Marin et al., 2002; Wong et al., 2002). There is also evidence that the development of the antennal lobe (AL), the first olfactory center, underlies stereotypic processes and is at least partly independent of sensory input. PNs target certain regions in the developing AL and form protoglomeruli before their presynaptic OSN partners arrive (Jefferis & Hummel, 2006; Jefferis et al., 2004). Even if the sensory input is eliminated directly after eclosion by removal of the olfactory organs, the antennae and maxillary palps, the morphology of PN axons and dendrites remains unaltered (Wong et al., 2002).

However, it has already been shown that at least some parts of the central nervous system of *D. melanogaster* are able to undergo experiencedependent plastic changes. Several studies demonstrated volumetric changes in different neuropils after flies perceived certain environmental stimuli such as low or high population densities (Heisenberg et al., 1995), or exposure to light (Barth et al., 1997) or to odors (Chodankar et al., 2020; Devaud et al., 2001; Devaud et al., 2003; Golovin et al., 2019; Kidd & Lieber, 2016; Kidd et al., 2015; Sachse et al., 2007). Similar effects could also be observed in other groups of arthropods, notably in Hymenoptera (Andrione et al., 2017; Arenas et al., 2012; Brown et al., 2004; Hagadorn et al., 2021; Hourcade et al., 2009; Jernigan et al., 2021; Penick et al., 2021; Withers et al., 1993), in Lepidoptera (Eriksson et al., 2019), and in spiders (Steinhoff et al., 2018).

Although it has been known for the past two decades that the volume of glomeruli in the olfactory system of *D. melanogaster* is affected by long-term exposure to odorants, the underlying cellular processes and cell types involved remained largely elusive so far (Fabian & Sachse, 2023). In our study, we focused on the highly specific olfactory circuit that is involved in the detection of the aversive odorant geosmin (Becher et al., 2010; Stensmyr et al., 2012). This odor ligand is solely detected by the Or56a receptor, which in turn is selectively tuned to geosmin. Moreover, as an indicator for the presence of potentially toxic microorganisms (Gerber & Lechevalier, 1965; Jüttner & Watson, 2007; Mattheis & Roberts, 1992), geosmin is considered to be a particularly important cue for the fly. In our study, we investigated the effects of geosmin exposure on the glomerular volume, the morphology of innervating neurons and glia cells, the signaling within the circuit and the behavior of flies. We show that even the highly specialized circuit that conveys information about the presence of potentially lethal threats underlies modulation by previous experience and enables the fly to adapt to a changing environment.

## RESULTS

### Long-term exposure to geosmin causes a volumetric increase of the DA2 glomerulus

Our first goal was to establish an exposure setup and procedure in our lab to replicate plasticity effects that were previously seen in other glomeruli after exposure to certain odorants (Chodankar et al., 2020; Devaud et al., 2001; Devaud et al., 2003; Golovin et al., 2019; Kidd & Lieber, 2016; Kidd et al., 2015; Sachse et al., 2007) in the geosmin-detecting glomerulus DA2. We exposed flies that express photoactivatable GFP in a subset of PNs (Fig. 1A) in an alternating manner of 1 min odor exposure and 5 min fresh air to geosmin during their early adult life (Fig. 1B) and subsequently measured the volume of the DA2 glomerulus. We could observe a significant volumetric increase of the DA2 glomerulus in both sexes after flies were exposed to geosmin (Fig. 1C). For the sake of longevity during photoactivation experiments and ecological relevance for behavior (e.g. oviposition), we focused on females in succeeding experiments for which we also controlled the glomerulus volume (Fig. S1, Table S1).

**Figure 1.**
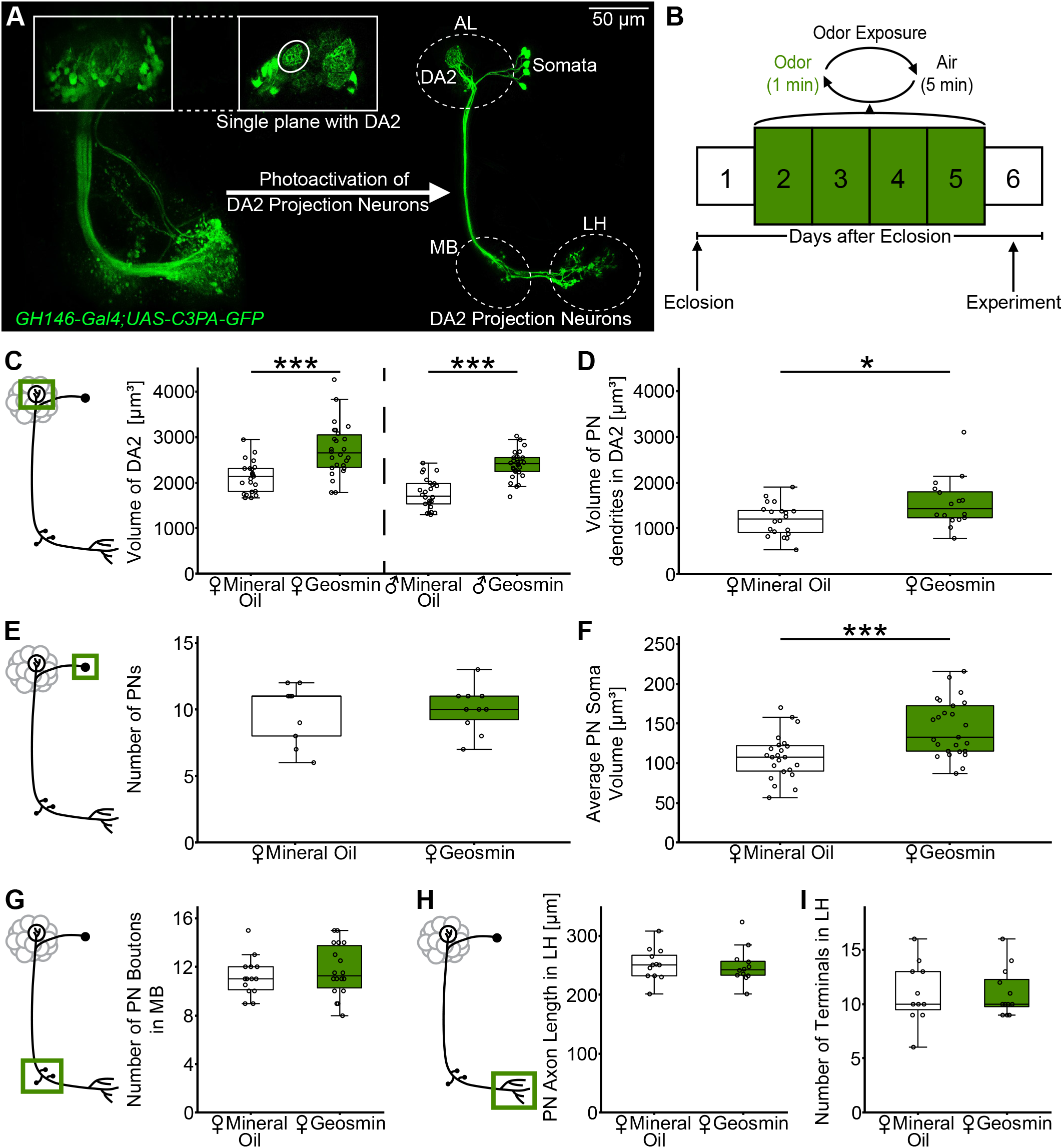
Long-term exposure to geosmin alters the morphology of DA2 projection neurons at the antennal lobe level but not in higher brain centers. **A**, schematic of the photoactivation approach showing representative z-stacks of the whole GH146-Gal4 population (left) and DA2 projection neurons (right). **B**, schematic of the odor exposure procedure used in this study. **C**, exposure to geosmin leads to a volumetric increase of the DA2 glomerulus in females and males (n = 23-28). **D**, exposure to geosmin increases the volume of the projection neurons dendrites within the DA2 glomerulus (n = 16-20). **E**, the total amount of projection neurons (summed up for both hemispheres) remains the same after geosmin exposure (n = 9-10). **F**, the average soma volume of projection neurons is increased in flies that were exposed to geosmin (n = 23-25). **G**, the number of projection neuron boutons in the mushroom body is unaltered after geosmin exposure (n = 14-18). **H, I**, the axonal length and axonal terminals of projection neurons in the lateral horn are not affected by the geosmin exposure (n = 11-12). Statistical test: two-sample student’s t test. Significance levels: * p < 0.05, *** p < 0.0005. Abbreviations: AL, antennal lobe; LH, lateral horn; MB, mushroom body, PN, projection neuron.

### DA2 projection neurons undergo plastic changes at the level of the antennal lobe but not in higher brain centers

To analyze the cellular basis causing the observed volumetric increase of the DA2 glomerulus, we first focused on second-order olfactory neurons, i.e. PNs. For this purpose, we quantified GFP positive voxels inside the DA2 glomerulus and observed that the volume of PN dendrites was significantly increased after geosmin exposure (Fig. 1D). According to this observation, we hypothesized that in geosmin exposed flies either the PNs extended their dendritic fields or that the DA2 glomeruli were innervated by a higher number of PNs. We quantified the number of PNs on both sides of the AL. However, we found that the number of PNs was not affected by exposure to geosmin (Fig. 1E), while the average size of PN somata was significantly increased (Fig. 1F). In addition to our findings in the region of the AL, we also evaluated whether the exposure had an effect on the PN morphology downstream of this neuropil. Counting PN boutons in the region of the mushroom body (MB) did not reveal any difference between the experimental and control group (Fig. 1G). Moreover, we reconstructed the terminal axonal branches in the LH, beginning at the site where axons enter the LH. We could only reconstruct one common skeleton tree for all PNs that innervate the DA2 glomerulus as we could not discriminate precisely between individual PNs in the region of the LH due to densely packed axons. However, it was still possible to quantify the overall length of axons and the number of their terminals. We did not find a difference in either of these two parameters after exposure to geosmin (Fig. 1H, I). Taken together, our data show that DA2 PNs from flies exposed to geosmin expand their dendrites and thereby contribute to the overall volume increase of the glomerulus, whereas their axonal terminals in the MB and LH remain unaffected.

As we could only reconstruct a common skeleton tree for DA2 PNs because they were so densely packed, we additionally exposed flies to 3-hexanone or ethyl hexanoate, ligands that strongly activate the olfactory circuits associated to the DM1 and DM2 glomeruli. Both glomeruli are only innervated by 1 or 2 PNs (Fig. S2A, F), which allowed reconstructing their terminals in the LH much more precisely. We could document a volumetric growth of both glomeruli (Fig. S2B, G), however, neither the number of boutons in the MB (Fig. S2C, H) nor the axon length (Fig. S2D, I) or number of axonal terminals (Fig. S2E, J) in the LH were affected. From these experiments, we conclude that the morphological changes of PNs are restricted to the region of the AL and do not occur in higher processing centers as the LH and the MB.

### Neurites of local interneurons are remodeled after long-term exposure to geosmin

Previous studies hypothesized that the glomeruli increase in size is due to increased branching and synaptic connectivity between PNs and local interneurons (LNs) (Das et al., 2011; Sachse et al., 2007). While we and others (Chodankar et al., 2020) found evidence that PNs indeed extend their dendritic fields, little is known about morphological plasticity effects in LNs. To elucidate whether the morphology of LNs is also affected by exposure to geosmin, we exposed flies that express photoactivatable GFP (C3PA-GFP) in different subpopulations of LNs by using enhancer-trap lines that label a diverse set of LNs, including some with a global as well as a patchy AL innervation (GH298, NP2426, NP3056, HB4-93) (Fig. 2A-D). We measured the volume of all LNs labeled by these different driver lines in the geosmin-responsive glomerulus DA2 as well as the DA4l as a control glomerulus by quantifying GFP positive voxels (Fig. 2A1-D1). In one of the LN lines (Fig. 2A1, GH298), we observed that the volume of LNs inside the DA2 glomerulus did indeed increase in geosmin-exposed flies compared to the control flies. This was distinct in the case of the DA2 glomerulus and could not be observed in the DA4l glomerulus. However, in the other LN lines we could not observe such an effect (Fig. 2B1-D1).

**Figure 2.**
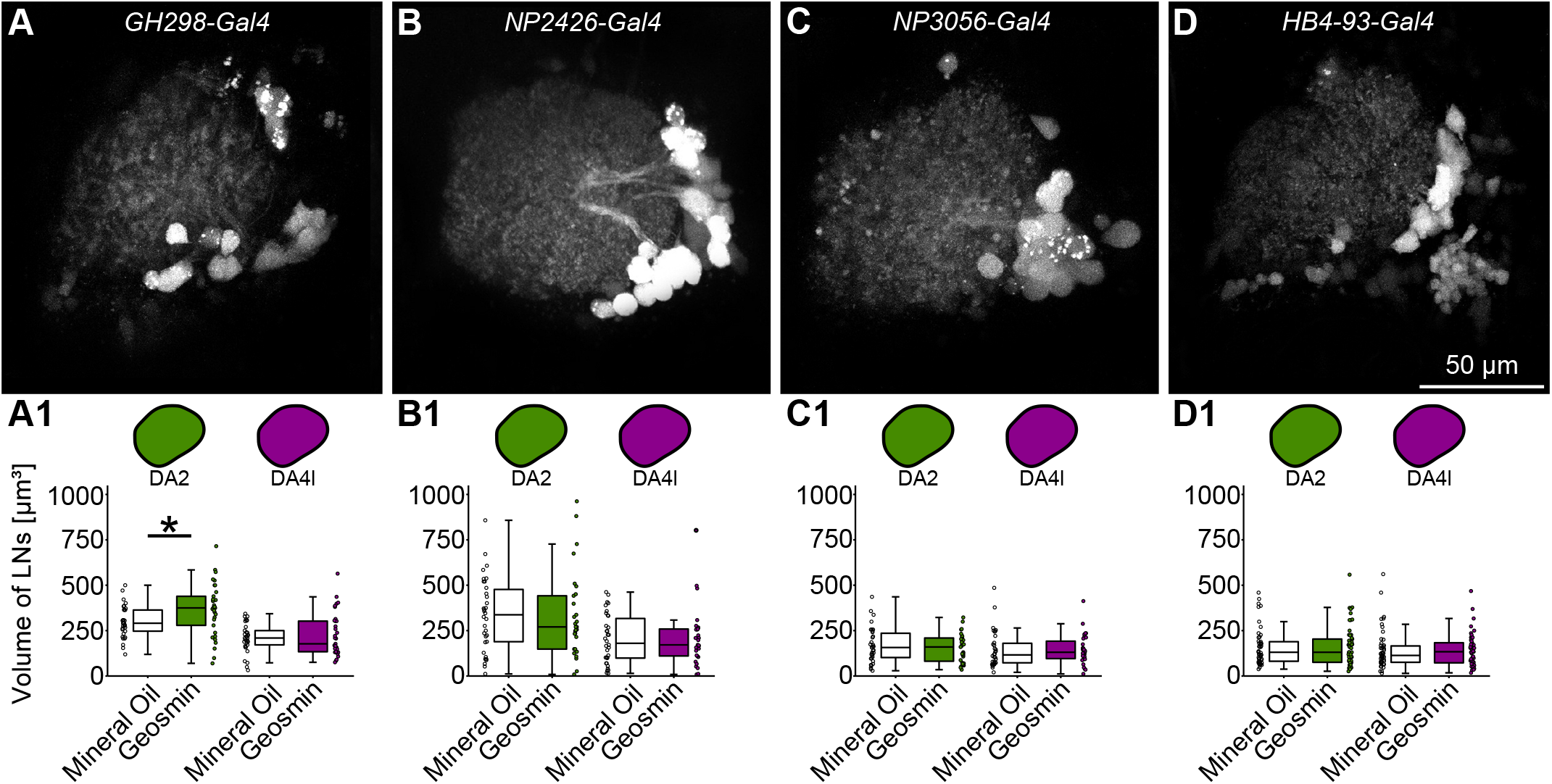
Morphological effects due to long-term exposure to geosmin can be seen in the entire local interneuron subpopulation of GH298-Gal4. **A-D**, representative z-projections of the different local interneuron lines used for the experiment. The expression of UAS-C3PA-GFP was driven by GH298-Gal4 (A), NP2426-Gal4 (B), NP3056-Gal4 (C) and HB4-93-Gal4 (D). A-C, global lines, D, patchy line. **A1-D1**, boxplots showing the volume that is occupied by the entire local interneuron subpopulation in the DA2 and DA4l glomeruli. The volume of GH298-Gal4 local interneurons (A1) in the DA2 glomerulus is increased due to exposure to geosmin, while the volume in the DA4l glomerulus is unaffected. This effect could not be seen in the other local interneuron lines. Sample sizes: A1-A3 (n = 33); B1-B3 (n = 29-34); C1-C3 (n = 31-34); D1-D3 (n = 50-53). Statistical test: two-sample student’s t test. Significance levels: * p < 0.05, *** p < 0.0005. Abbreviation: LNs, local interneurons.

In order to pinpoint specific cellular morphological plasticity effects in GH298 LNs, we used photoactivation of photoactivatable GFP to label single LNs. We targeted single LN somata during the photoactivation procedure and reconstructed those parts of the LN that innervated the DA2 glomerulus (Fig. 3A) and a second glomerulus (DA4l/DC1), which should not be affected by exposure to geosmin. We consistently selected a cell body in the same location in different flies to reduce heterogeneity in our samples. Three of the LN lines (GH298, NP2426 and NP3056) label mostly global LNs, which innervate almost all glomeruli in the AL (Fig. 3B-D), while the fourth LN line (HB4-93) labels a variety of different LN subtypes. In the fourth line we focused on patchy LNs (Chou et al., 2010) that innervate approximately half of the glomeruli of the AL (Fig. 3E). We chose the DC1 glomerulus as our control glomerulus instead of the DA4l in this line, as we selectively labeled individual LNs that innervated both the DA2 and DC1 more frequently than LNs that innervated the DA2 and DA4l. The total length of the LN branches inside the glomeruli (Fig. 3B1-E1) and the number of the LN terminals (Fig. 3B2-E2) were not affected by the exposure to geosmin. Although we did not observe any change in the number of terminals, we still found that the average distance between terminals significantly increased in NP3056 LNs in geosmin exposed flies (Fig. 3D3). We also quantified the number of boutons manually along the LN branches and divided it by the total length of the LN inside the glomerulus to obtain a measure of the bouton density (Fig. 3B4-E4). Notably, in the NP2426-Gal4 line, we observed an increase of the bouton density in the DA2 glomerulus after geosmin exposure, while this effect was absent in our internal control glomerulus DA4l (Fig. 3C4). Additionally, we measured the average volume of boutons (Fig. 3B5-E5) and found a significant increase in the GH298-Gal4 line, which was again specific for the DA2 glomerulus and could not be observed in the DA4l glomerulus (Fig. 3B5).

**Figure 3.**
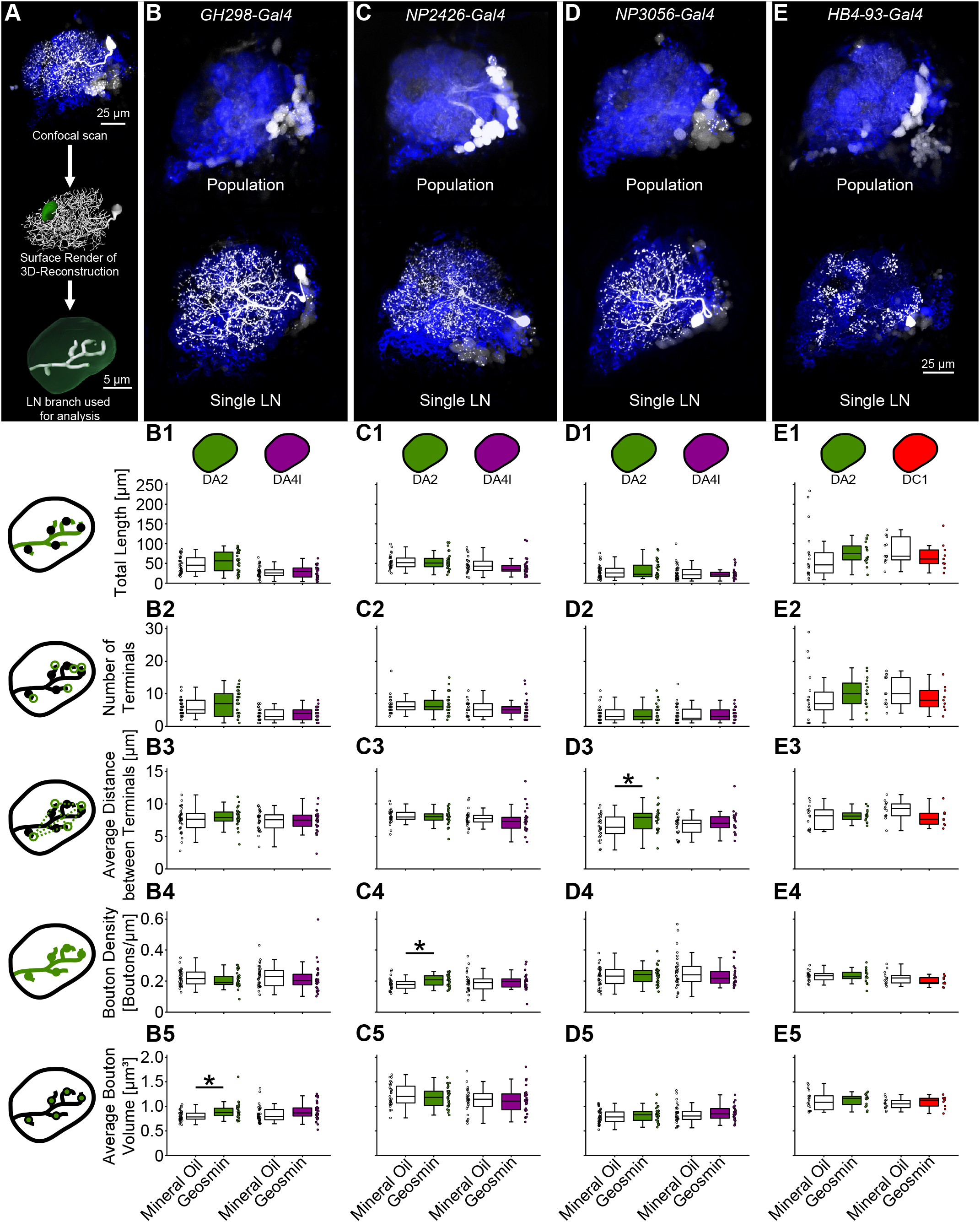
Long-term exposure to geosmin causes morphological effects in global local interneurons. **A**, schematic showing the acquisition of morphological data. In each fly a single local interneuron soma was photoactivated. A z-scan was acquired and the local interneuron part innervating the DA2 glomerulus (and control glomerulus DA4l/DC1) was reconstructed. The morphological parameters of the reconstruction were then analyzed. **B-E**, representative z-projections of the different local interneuron lines used for the experiment. The whole population (upper panel) and single photoactivated local interneurons (lower panel) are shown. B-D, global LNs, E, patchy LNs. **B1-E1**, the total length of the local interneuron inside of the glomerulus was unchanged by geosmin exposure. The icons in the top represent the experimental glomerulus (DA2) and the internal control glomeruli (DA4l/DC1). **B2-E2**, the number of local interneuron terminals is unaffected by exposure to geosmin. **B3-E3**, in the NP3056-Gal4 line (D3), the average Euclidean distance between terminals was significantly higher in geosmin-exposed flies than it was in the control group. The effect was specific to the DA2 glomerulus and could not be seen in the control glomerulus DA4l. **B4-E4**, in the NP2426-Gal4 line (C4), the bouton density increased due to the exposure to geosmin. The effect was specific to the DA2 glomerulus and could not be seen in the control glomerulus DA4l. **B5-E5**, in the GH298-Gal4 line (B5), the average bouton volume was significantly higher in geosmin exposed flies than it was in the control group. The effect was specific to the DA2 glomerulus and could not be seen in the control glomerulus DA4l. Sample sizes: B1-B5 (n = 26-33); C1-C5 (n = 26-29); D1-D5 (n = 20-35); E1-E5 (n = 9-18). Individual data points are indicated as scatterplots next to the respective boxplots. Statistical test: two-sample student’s t test. Significance levels: * p < 0.05. Abbreviation: LN, local interneuron.

Boutons are neuronal structures, which indicate the presence of synapses (Rybak et al., 2016). Therefore, both observed plasticity effects of LN boutons in our single cell experiments (Fig. 3B5, C4) could be an indicator that the total number of synapses of LNs increased due to the exposure to geosmin. Consequently, we evaluated whether there are similar effects when we label synapses of entire LN populations in our target glomeruli. For the quantification of LN synapses, we used flies expressing GFP tagged to Brp-Short as a presynaptic marker in the NP2426-Gal4 and GH298-Gal4 lines (Fig. 4A). Although both LN lines label mostly global LNs, which innervate the majority of glomeruli in the AL, the distribution of Brp-Short puncta across the entire AL differs strikingly between the two lines (Fig. 4B, F). In the NP2426-Gal4 line, Brp-Short puncta were distributed evenly and the Brp-Short signal alone was sufficient to identify some glomeruli (Fig. 4B). However, in the GH298-Gal4 line, Brp-Short puncta were scattered more irregular and the signal consisted of regions with dense clusters and regions with only very few puncta (Fig. 4F). Glomerulus borders were much more ambiguous in the GH298-Gal4 line. We could not observe a change in the number of Brp-Short puncta in the DA2 or DA4l glomerulus in these two lines (Fig. 4C, G). However, the average volume of Brp-Short puncta decreased in both investigated glomeruli in the NP2426-Gal4 line after geosmin exposure (Fig. 4D). We could observe the same phenomenon in the GH298-Gal4 line, but the effect was specific to the DA2 glomerulus and could not be observed in the DA4l glomerulus (Fig. 4H). When we quantified the fluorescence intensity of the DA2 and DA4l glomeruli in our initial confocal scans, we found that the relative fluorescence of the DA2 glomerulus decreased in both lines after exposure to geosmin (Fig. 4E, I), indicating that the observed decreased average volume of Brp-Short puncta was not artificially introduced by deconvolution or 3D watershedding.

**Figure 4.**
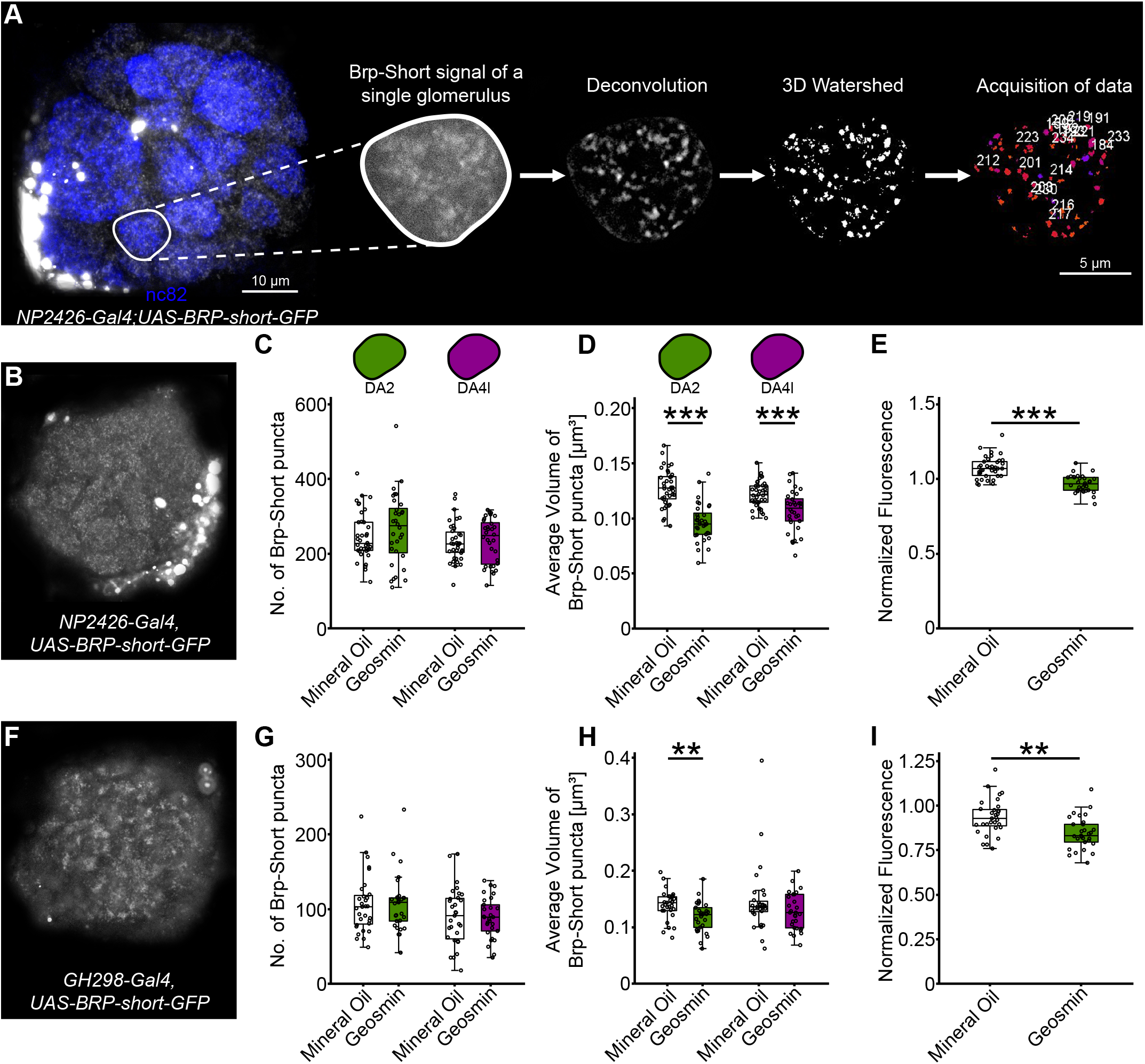
Long-term exposure to geosmin has an impact on the morphology of LN presynapses in the DA2 glomerulus. **A**, schematic of the process of quantifying LN presynapses in the DA2 glomerulus. We scanned immunostained brains of flies expressing GFP tagged to Brp-Short. The GFP signal of individual glomeruli was extracted and deconvolved. 3D watershedding split clusters of Brp-Short puncta into individual structures, which were later counted and measured automatically. **B**, exemplary image of the Brp-Short-GFP signal in the NP2426-Gal4 line. The image shows a single plane in the middle of the AL. **C**, the numbers of Brp-Short puncta in the DA2 and DA4l glomeruli were not affected by exposure to geosmin. **D**, the average size of Brp-Short puncta was reduced in both glomeruli after exposure to geosmin. **E**, the normalized fluorescence signal of the DA2 glomerulus decreased due to exposure to geosmin. **F**, exemplary image of the Brp-short-GFP signal in the GH298-Gal4 line showing a single plane in the middle of the AL. **G**, the numbers of Brp-Short puncta in the DA2 and DA4l glomeruli were not affected by exposure to geosmin. **H**, the average size of Brp-Short puncta was reduced in the DA2 glomerulus but not in the DA4l glomerulus after exposure to geosmin. **I**, the normalized fluorescence signal of the DA2 glomerulus decreased due to exposure to geosmin. Statistical test: two-sample student’s t test. Sample sizes: n = 29-38. Significance levels: ** p < 0.005, *** p < 0.0005. Abbreviation: LN, local interneuron.

In summary, we found that the GH298 LN subpopulation contributes slightly to the growth of glomerulus DA2 after exposure to geosmin. Photoactivation of individual LNs revealed subpopulation-specific effects at the level of LN terminals and boutons, some of which indicate synaptic changes. In addition, quantifications of Brp-Short puncta confirmed that LN synapses are affected. These experiments demonstrate that different LN subpopulations undergo different morphological changes during longterm exposure to geosmin.

### OSNs and glia cells do not contribute to the volumetric increase in the DA2 glomerulus

We also evaluated whether OSNs contribute to the volumetric effect that occurs in the DA2 glomerulus after exposure to geosmin or whether they retract their axons as was recently shown for the VM7d glomerulus (Chodankar et al., 2020; Golovin et al., 2019). We analyzed the volume of OSNs by quantifying GFP positive voxels in the DA2 glomerulus of flies that express GFP under the control of the Orco-Gal4 driver line (Fig. 5A). We did not observe any change in absolute OSN axon volume in the DA2 glomerulus (Fig. 5B). Due to the long-term exposure to geosmin, Or56a OSNs are constantly activated leading to an increased energy consumption that requires efficient Ca^2+^ buffering (Berridge, 1998). We therefore hypothesized that these neurons might upregulate their metabolism, which is known to affect the morphology of mitochondria (Westermann, 2012). To verify that, we used flies that express GFP tagged to mitochondria in Or56a OSNs, which we quantified in a similar way as described for the quantification of Brp-Short puncta (Fig. 5C). Although the total volume of mitochondria in Or56a OSNs in the DA2 glomerulus was not affected by exposure to geosmin (Fig. 5D), we observed a significant increase in the total number of mitochondria (Fig. 5E). This increase was accompanied by a lower average volume of mitochondria (Fig. 5F), indicating a mitochondrial fission process.

**Figure 5.**
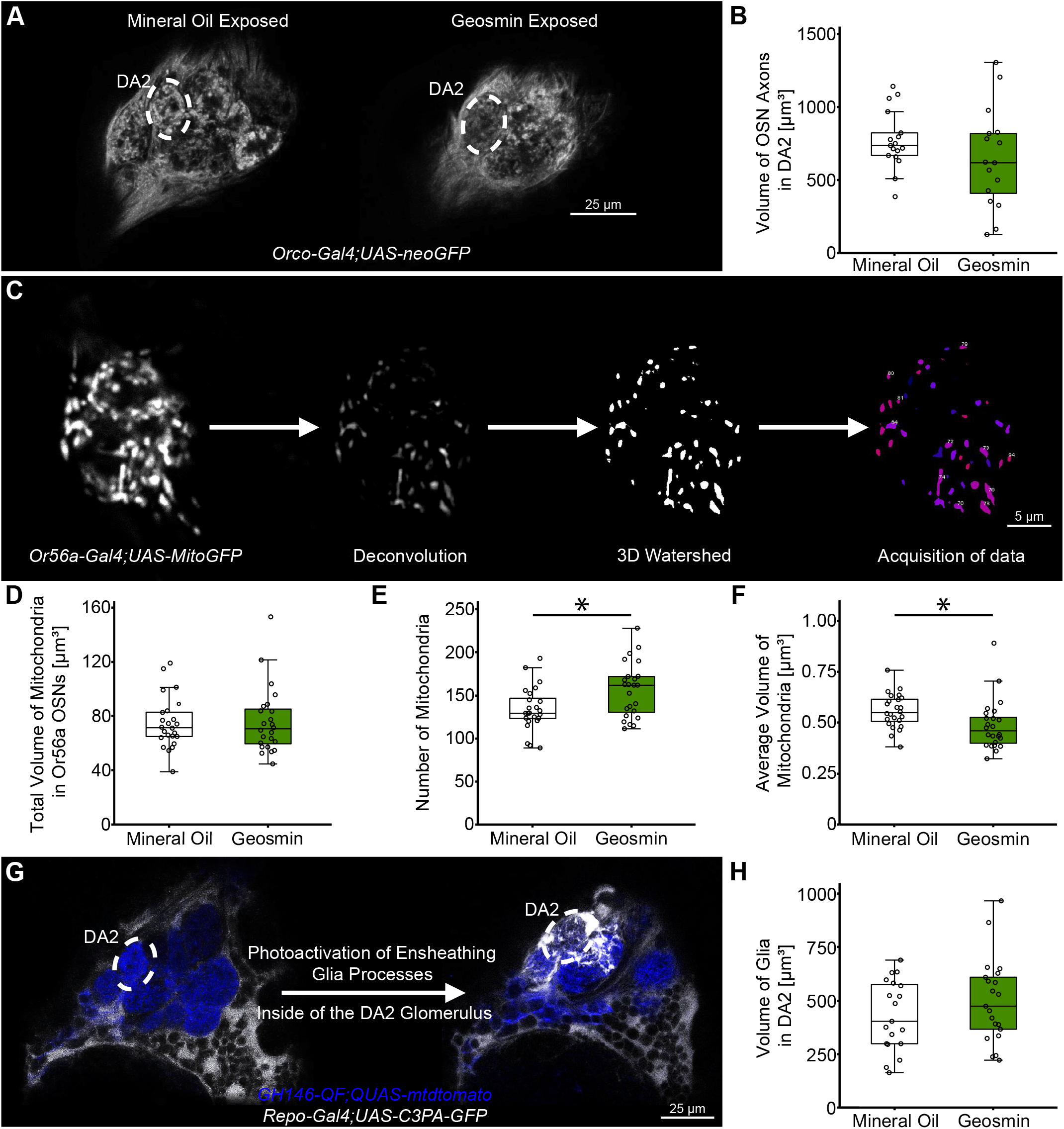
Effects of long-term exposure on olfactory sensory neurons and glia cells. **A**, example images showing the neoGFP signal in OSNs of mineral oil (left) and geosmin (right) exposed flies. **B**, the volume of OSN axons innervating the DA2 glomerulus is not significantly affected by exposure to geosmin (n = 16-17). **C**, schematic of the process of quantifying mitochondria in Or56a OSNs. Confocal image stacks were cropped and deconvoluted. 3D watershedding was used to split clusters of several mitochondria in individual structures. The mitochondria were then counted and their volume was measured automatically. **D**, the total volume of mitochondria in Or56a OSNs in the region of the DA2 glomerulus was not affected by exposure to geosmin (n = 24). **E**, the amount of mitochondria in Or56a OSNs in the region of the DA2 glomerulus increased due to exposure to geosmin (n = 24). **F**, the average volume of mitochondria in Or56a OSNs in the region of the DA2 glomerulus decreased due to exposure to geosmin (n = 24). **G**, scheme of the photoactivation process of ensheathing glia cells of the Repo-Gal4 line. Only ensheathing glia cells that send processes into the DA2 glomerulus were photoactivated. **H**, the volume of ensheathing glia cell processes in the DA2 is not significantly affected by exposure to geosmin (n = 19-21). Statistical test: two-sample student’s t test. Significance levels: * p < 0.05. Abbreviation: OSN, olfactory sensory neuron.

To clarify whether non-neuronal cells also contribute to the glomerular volume increase due to long-term geosmin exposure, we analyzed glia cells. In the AL, glia cells are mainly located between glomeruli and form a sheath around them. Some of them extend cell processes into glomeruli and could therefore contribute to the volumetric increase of the DA2 glomerulus after exposure to geosmin. We used flies that express photoactivatable GFP in the pan-glial Repo-Gal4 driver line (Sepp et al., 2001). To measure the volume of labeled glia cell processes, we photoactivated glia cells in the DA2 glomerulus (Fig. 5G) and quantified GFP positive voxels. As the exposure to geosmin had no effect on the volume of glia cells within the DA2 glomerulus (Fig. 5H), we conclude that they do not contribute to the glomerular growth.

### Long-term exposure to geosmin alters behavioral preference

A previous study showed that geosmin has an innate repellent effect on *D. melanogaster* in different behavioral assays (Stensmyr et al., 2012). We evaluated whether long-term exposure indeed affects the behavioral response towards this odorant. Exposed females were tested for their general approach response in a T-maze assay (Fig. 6A). We observed that flies previously exposed to mineral oil were significantly repelled by geosmin, while flies previously exposed to geosmin exhibited a neutral response (Fig. 6B). Since the odor exposure experiments were performed with fly vials containing food, the flies were always exposed to food odor in addition to geosmin. We therefore tested whether the context of the odor presentation has an impact on the behavior by adding a small cube of fly food on top of the filter paper. Again we observed that flies exposed to mineral oil were significantly repelled by geosmin, while the choice of geosmin-exposed flies was indifferent (Fig. 6C). Since geosmin-exposed flies were not generally repelled by the odorant anymore, we tested whether they also accept a source of geosmin as their oviposition substrate (Fig. 6D). Similar to the results of the T-maze assays, control flies exposed to mineral oil clearly avoided the substrate containing geosmin for oviposition, while geosmin-exposed flies did not exhibit a preference (Fig. 6E). These results show that long-term exposure to geosmin diminishes the repellent effect of this odorant.

**Figure 6.**
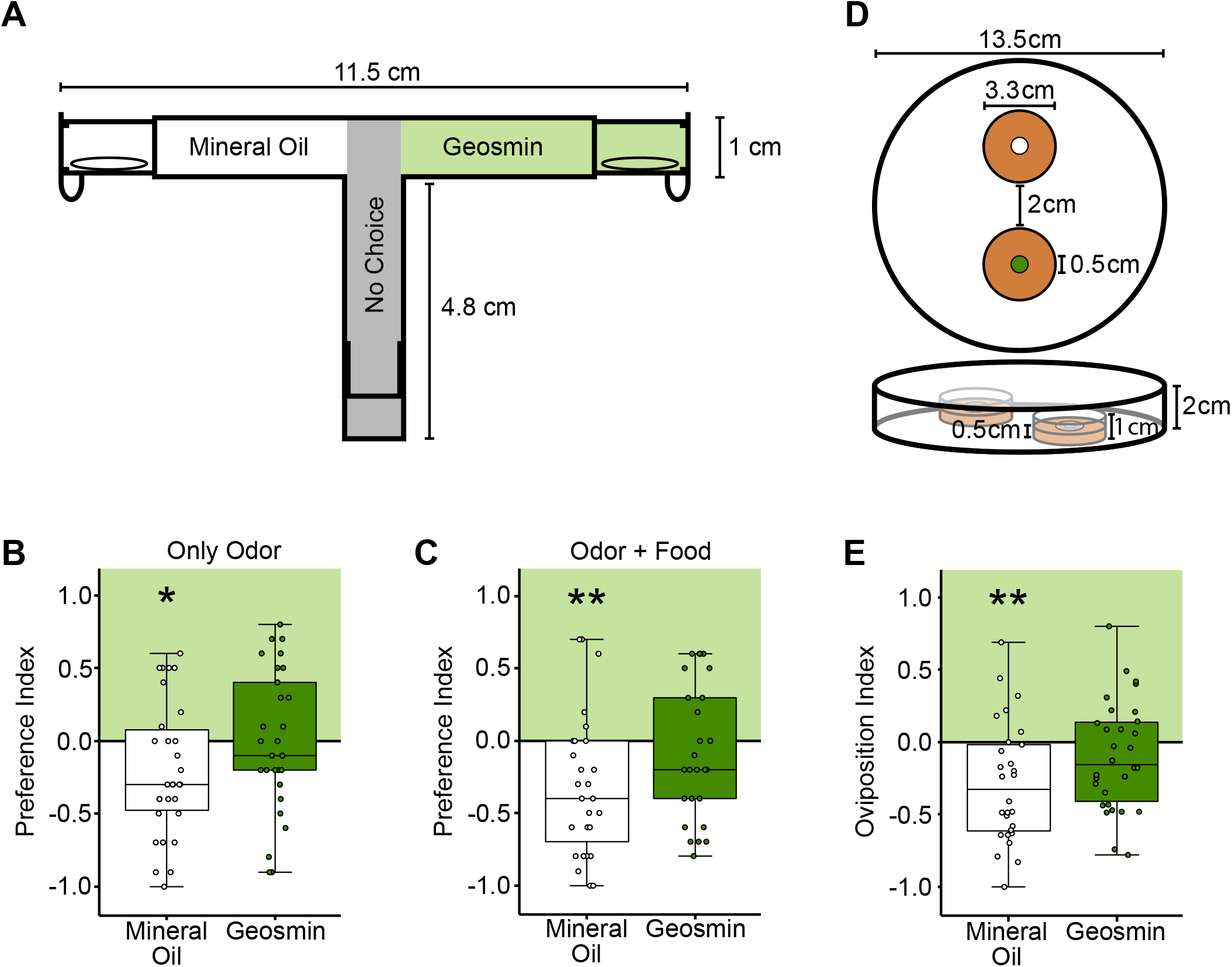
Long-term exposure to geosmin reduces aversion in *D. melanogaster* females. **A**, schematic of the T-maze setup. **B-C**, geosmin-exposed flies behave indifferently to geosmin, while mineral oil-exposed flies are repelled by the odor, independent of whether only the odor (B) or the odor with food (C) is presented. Sample size: n = 26-30. **D**, schematic of the oviposition assay. **E**, geosmin-exposed flies did not show a preference for the oviposition substrate, while mineral oil-exposed flies were repelled by the odorant. Sample size: n = 28-30. Error bars in bar plots indicate the standard error of mean. Statistical test: one-sample student’s t test. Significance levels: * p < 0.05, ** p < 0.005.

### Long-term exposure to geosmin causes no physiological effects in the underlying circuit

Having observed that flies exposed to geosmin tolerate this odorant, we tested whether the underlying signaling between neurons in this particular circuit is modified by the exposure. We hypothesized that the PN output response in geosmin-exposed flies is reduced by lateral inhibition by LNs, which could lead to less aversion towards the odor as previously shown for the repellent CO2 circuit (Sachse et al., 2007). This previous study did not show any modulation at the OSN level. To verify that the OSN output signaling is unaffected in the geosmin circuit, we employed flies expressing GCaMP3 coupled to synaptophysin (syp), a presynaptic vesicle protein (Pech et al., 2015), in these neurons. We delivered geosmin in five different concentrations (10^−7^-10^−3^) to the antenna of dissected flies and monitored odor-evoked calcium responses in the AL. The dose response curves of presynaptic OSN responses (Fig. 7A) showed an increase up to a geosmin concentration of 10^−4^, with a drastic drop in signal intensity at 10^−3^. As expected, we could not observe any significant difference between both treatments.

**Figure 7.**
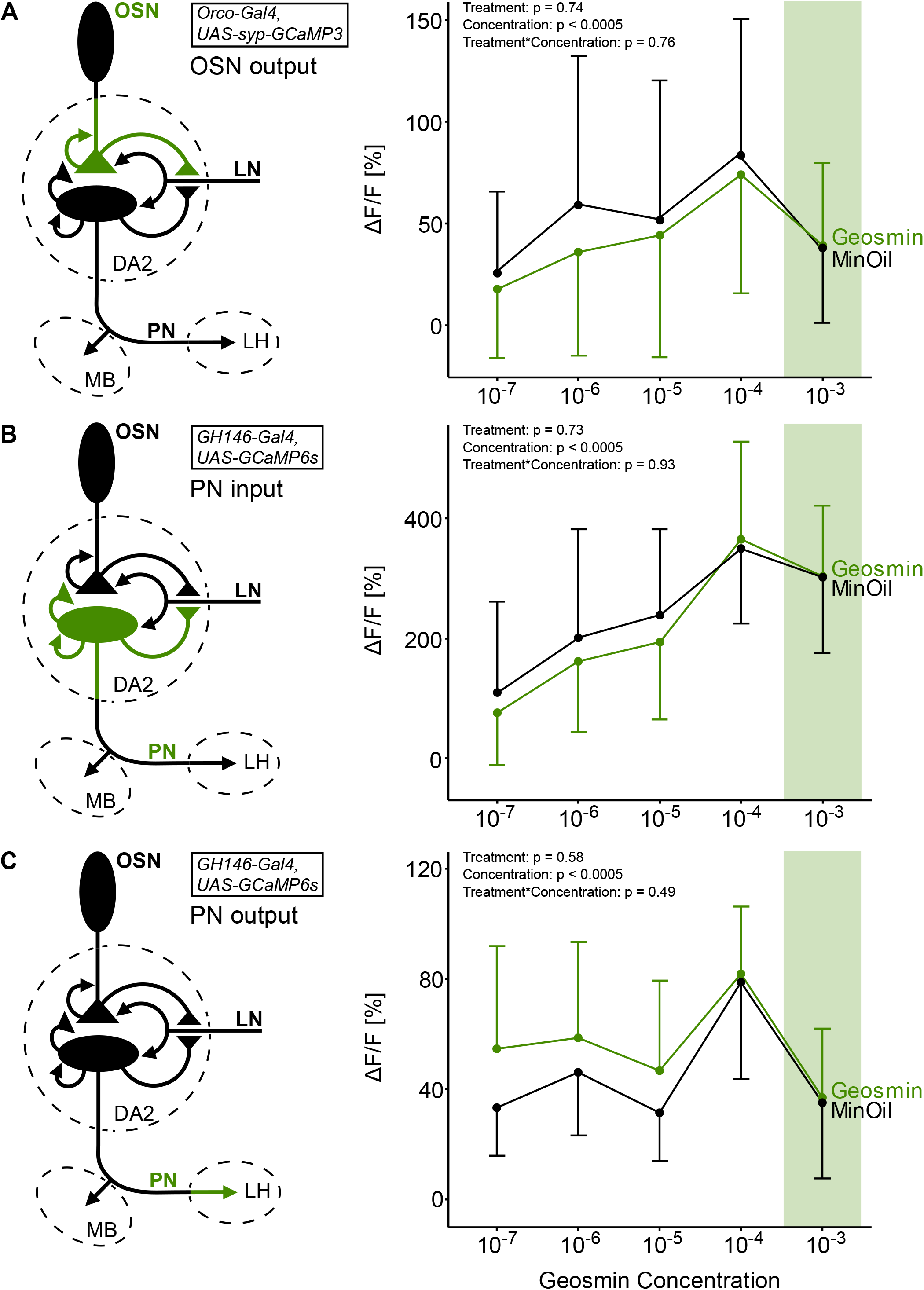
Calcium imaging responses to geosmin in different neurons of the DA2 glomerulus. **A**, presynaptic signaling in DA2 OSNs was not affected by exposure to geosmin (n = 12). **B**, signaling in DA2 PNs was not affected by exposure to geosmin (n = 12-13). **C**, signaling of DA2 PNs in the LH is increased for low geosmin concentrations when flies were previously exposed to geosmin by exposure to geosmin (n = 17). Error bars: standard deviation of mean. Dose response curves were tested with a two-way repeated measures ANOVA and Tukey’s post-hoc test. The schematics show the neuron type and the part of the neurons that was measured (green). Green rectangles show the geosmin concentration that flies were previously exposed to. Abbreviations: LH, lateral horn; LN, local interneuron; MB, mushroom body; OSN, olfactory sensory neuron; PN, projection neuron.

To monitor geosmin-evoked PN responses, we used flies that express UAS-GCaMP6s driven by the GH146-Gal4 driver. Because PNs receive input mostly in the AL and give the majority of their output in the MB and LH, input and output signals of PNs can be quantified by measuring their responses in the respective neuropil. In a first step, we measured PN input responses in the DA2 glomerulus (Fig. 7B). The dose response curves closely resembled those observed for the presynaptic odor responses in OSNs (Fig. 7A), without a significant difference between the treatments. The measurements in the LH again revealed a concentration-dependent increase up to 10^−4^ with a subsequent decrease at 10^−3^ (Fig. 7C). The only difference we observed in the general trend of the curves was a lower response for the 10^−5^ concentration. The dose response curves of both treatment groups were not significantly different from each other. Geosmin-exposed flies showed a slightly stronger response at lower geosmin concentrations. As a whole, the results of the physiology experiments suggest that the exposure to geosmin does not significantly alter the signal transduction in the olfactory circuit.

## DISCUSSION

### Morphological changes due to long-term odor exposure

Our data show that one of the most specific circuits in the fly’s olfactory system, which conveys information about the presence of potentially lethal microorganisms (Stensmyr et al., 2012), is subject to distinct structural changes that depend on previous experience. This includes an increased glomerular volume, an expansion of PN dendrites, morphological changes at the level of LN synapses, and even subcellular changes like the fission of mitochondria in OSNs. These transformations were accompanied by behavioral changes, with geosmin-exposed flies tolerating the otherwise aversive odorant, while the neuronal responses within the circuit, comprising first- and second-order neurons, were mostly unaffected.

Our experiments consistently revealed a specific growth of the DA2 glomerulus when flies where exposed to geosmin for four consecutive days. Additionally, we observed the same effect in two other glomeruli with odorants that were not investigated in this context before (DM1 with 3-hexanone and DM2 with ethyl hexanoate), which suggests that the glomerular growth in response to extensive odorant exposure is a general phenomenon in the AL of *D. melanogaster*. However, very early studies on this phenomenon reported a decrease in glomerular vol-ume after long-term odorant exposure (Devaud et al., 2001; Devaud et al., 2003), while a recent study showed that the volume of the VM7d glomerulus is unaffected in flies exposed to ethyl butyrate (Chodankar et al., 2020). The limited data on volumetric changes caused by long-term odorant exposure indicated a general trend that glomeruli grow in size when the associated circuit is stimulated by a strong ligand. However, some glomeruli react differently to this stimulation by showing a decreased volume or no volumetric change at all.

Although the phenomenon of glomerular growth is known for more than a decade (Chodankar et al., 2020; Das et al., 2011; Kidd & Lieber, 2016; Kidd et al., 2015; Sachse et al., 2007), the underlying cellular processes are still largely elusive. We evaluated which neurons contribute to the volumetric changes, and hypothesized that the PNs innervating the DA2 glomerulus could be responsible, as their number is very variable. However, our data demonstrate that the number of PNs innervating the DA2 glomerulus is not changed after exposure, whereas their dendrites expand. This finding is conform with the results of a recent study showing a volumetric increase in PN dendrites of the DM5 and V glomeruli after flies were exposed to ethyl butyrate or CO2, respectively (Chodankar et al., 2020). Additionally, we observed a significant increase of the average soma volume of DA2 PNs that could be related to the extension of PN dendrites. The larger dendritic field is an indicator that the metabolic and energetic requirements of the PNs could be increased. In fact, a study on crabs (Brachyura) showed that the soma size of neurons correlates with the expression of glyceraldehyde-3-phosphate dehydrogenase (GAPDH), 18S rRNA and EF1-alpha (Ransdell et al., 2010). These gene products are involved in basal metabolic cell functions. GAPDH, for instance, is an important enzyme involved in the energy metabolism of cells (Sirover, 1999). Therefore, a larger PN soma could potentially compensate for additional metabolic requirements.

These observations lead to the question why PNs extend their dendrites. One explanation could be that this increases the space for additional synaptic contacts. A study on photoreceptor neurons in the housefly Musca domestica demonstrated that membrane area and number of synapses are strictly proportional (Nicol & Meinertzhagen, 1982). It is conceivable that such a correlation also occurs in PNs in the olfactory system of *D. melanogaster*. Tobin et al. (2017) showed that the synaptic density between OSNs and individual PNs is very similar between sister PNs of the DM6 glomerulus in both brain hemispheres, even when they differ strikingly in their length. If the synapse density is also independent of the length in DA2 PNs, we can expect that they form more synapses when they extend their dendritic fields. Recent connectome studies showed unambiguously that PNs mainly get input from OSNs and LNs (Horne et al., 2018; Schlegel et al., 2021; Tobin et al., 2017). LNs are a morphologically very diverse group of AL neurons (Chou et al., 2010; Tanaka et al., 2012) that can either be inhibitory (Olsen & Wilson, 2008; Root et al., 2008; Wilson & Laurent, 2005) or excitatory (Assisi et al., 2012; Huang et al., 2010), which renders them also functionally diverse. We hypothesized possible morphological effects in the LNs as potential synaptic partners for PNs and therefore quantified the LN branches of entire LN lines, as well as the branches of individual LNs innervating the DA2 glomerulus. From our four LN lines investigated (i.e. GH298, NP2426 and NP3056 labeling global LNs; HB4-93 labeling diverse LNs), we observed significant effects In two global interneuron lines; we found that the boutons are affected by the exposure in the form of an increased bouton density in the NP2426-Gal4 line and an increased average bouton volume in the GH298-Gal4 line. Since boutons are potential regions for synaptic contacts between neurons (Rybak et al., 2016), we hypothesized that these two effects could indicate that LNs of these two lines modulate the number or size of synapses. We tested this by quantifying Brp-Short puncta as a proxy for synapses (Mosca & Luo, 2014) in the NP2426-Gal4 and the GH298-Gal4 line. Brp is involved in the formation of the T-bar structure at presynaptic sites (Wagh et al., 2006) and is therefore a useful indicator for the presence of presynapses. Contrary to our expectations, we could not observe a quantitative change in Brp-Short puncta in either of the two LN lines. However, the average Brp-Short puncta volume decreased significantly in both lines indicating smaller synapses. This effect occurred in the DA2 glomerulus in the GH298-Gal4 line but not in the NP2426-Gal4 line. For the visual system of M. domestica, it is reasoned that a small synapse indicates that it was relatively newly formed (Fröhlich & Meinertzhagen, 1982; Rybak & Meinertzhagen, 1997). During development, these newly formed synapses sequentially add postsynaptic partners to a presynaptic site until they establish the mature tetrad configuration (Fröhlich & Meinertzhagen, 1983). It is conceivable that a similar process occurs in *D. melanogaster* LNs since they also form polyadic synapses that usually have a triad or tetrad configuration (Horne et al., 2018; Rybak et al., 2016). The results of the single cell photoactivation experiments and Brp-Short experiments are not necessarily contradicting each other. In our photoactivation experiments, we photoactivated LN somata in the same location to target homogeneous LNs in different flies, thus minimizing variability between individuals. The individual LNs we labeled could be affected in a specific way, whereas the rest of the LN subpopulation could be affected differently. This could obscure the changes observed in single-cell photoactivation experiments when the entire subpopulation is examined. The various morphological effects that we observed in different global LN lines in our single-cell photoactivation experiments support the idea that different subpopulations of LNs respond differently to long-term odor exposure. Chou et al. (2010) showed that the GH298-Gal4 and NP3056-Gal4 lines label largely non-overlapping LNs, but such data on NP2426-Gal4 LNs are presently unavailable. However, our Brp-Short staining experiment revealed that the distribution of Brp-Short puncta across the entire AL looks very different between the GH298-Gal4 and NP2426-Gal4 lines. Therefore, we assume that these two LN lines also label largely non-overlapping subpopulations of LNs, suggesting LN-specific effects. Our findings demonstrate that long-term exposure to geosmin causes complex and various structural changes in different LN subpopulations in the DA2 glomerulus.

As another potential synaptic partner for PNs, we examined the axonal terminals of OSNs that innervate the DA2 glomerulus. Since quantification of the volume of OSN axons did not reveal any difference between both treatment groups, we did not expect that they form more synaptic contacts with PNs. This observation is in agreement with observations in the CO2-detecting circuit (Sachse et al., 2007). However, contradictory results regarding the exposure effects on OSNs were recently published (Chodankar et al., 2020; Golovin et al., 2019). Both studies demonstrate that the OSNs innervating the VM7d glomerulus retract their axonal terminals distinctly after exposure to ethyl butyrate, suggesting a glomerulus-specific effect for OSN axons. Interestingly, even though we did not see a volume effect in OSN axons, we found that the mitochondria in DA2 OSNs were affected by geosmin exposure. Their number increased and they became smaller, whereas the total volume of all mitochondria in the axons of DA2 OSNs did not change. Oxidative stress is a factor that facilitates the fragmentation of mitochondria (Wu et al., 2011). It is conceivable that neurons in the geosmin circuit are subject to increased oxidative stress during the exposure period as the involved neurons are frequently stimulated. However, frequent stimulation is associated with an increased metabolic activity and a higher energy demand, which leads to fusion of mitochondria (Westermann, 2012) instead of fission. Since mitochondria sequester Ca^2+^ from the cytosol (Murphy & Steenbergen, 2021), it is possible that the increased surface area resulting from fission facilitates and accelerates Ca^2+^ uptake, which could reduce the risk of excitotoxicity. Despite of this ambiguity, our observation shows that long-term exposure to odorants cannot only affect cellular morphology in some neurons, but also organelles in neurons that remain otherwise unmodified.

In summary, our data show that the extension of PN dendrites contributes mainly to the volume of the DA2 glomerulus and also LNs, as potential synaptic partners of PNs, apparently contribute and are rearranged during the exposure period, whereas OSNs and glia cells seem to play a subordinate role. However, that we did not observe any morphological effects in OSN axons, glia cells or PN axons in the MB and LH does not necessarily mean that there are no changes. A study on developing neurons in the larval visual system of *D. melanogaster* showed that neurite terminals extend and retract to the same extent. Consequently, the total length does not change (Sheng et al., 2018), even though intensive dynamic processes occur. Such dynamic plasticity effects could only be revealed by observing the glomerular growth in real-time.

### Long-term exposure to geosmin elicits behavioral changes

Although some previous studies demonstrated that long-term exposure of odorants causes behavioral changes in flies (Chodankar et al., 2020; Das et al., 2011; Devaud et al., 2001; Devaud et al., 2003; McCann et al., 2011; Sachse et al., 2007; Sudhakaran et al., 2013), it was never shown that such changes can also be induced by a highly specific and unambiguous odorant like geosmin. Our T-maze and oviposition experiments demonstrated that female flies previously exposed to geosmin were subsequently less repelled by this odorant, likely due to an associative learning process (Dylla et al., 2023). This challenges the postulated predictability in the behavioral outcome of “labeled line” signaling in the olfactory system (Keesey & Hansson, 2021). Our results clearly show that the valence of an odorant that activates one of the most specific olfactory circuits can be altered by previous experience.

To elucidate the connection between our morphological and behavioral results, we conducted physiological experiments in different regions of the geosmin circuit. There was no effect on presynaptic OSN signaling (OSN output) after geosmin exposure. Consistent with this, PN input was also unaffected. Only the physiological measurements of DA2 PNs in the lateral horn (PN output) revealed a slight trend towards enhanced responses when geosmin-exposed flies were tested with low concentrations of this odorant. Our findings contradict some earlier results that either showed a general decrease (Das et al., 2011; McCann et al., 2011; Sachse et al., 2007; Sudhakaran et al., 2013) or an increase (Kidd & Lieber, 2016; Kidd et al., 2015) of PN signaling after long-term odor exposure. The consistency of the neuronal signals we observed within the circuit indicates that physiological changes downstream of PNs may be responsible for the behavioral effects, as also suggested by two recent studies (Dylla et al., 2023; Gugel et al., 2023) that obtained similar physiological results. It is conceivable that glomerulus- or odor-specific mechanisms cause the different physiological effects observed across studies. Another relevant factor could be the method of odor delivery, as both Gugel et al. (2022) and we used a pulsed exposure system, in contrast to the permanent exposure used in other studies. Our data suggest that it may be advantageous for the fly to keep signaling within the olfactory circuit constant when the involved neurons detect an odorant that represents a threat signal. How the responses remain largely unaffected despite the structural changes we observed remains unclear. However, it is conceivable that compensatory effects between different types of neurons in the glomerulus may be responsible.

Highly specific receptors such as Or56a or pheromone receptors are essential for the behavior as they translate crucial environmental information into neuronal signals. At first glance it appears counter-intuitive that the geosmin circuit that conveys information about the presence of toxic threats (Stensmyr et al., 2012) changes due to previous experience. However, plastic effects in neuronal tissue are an effective way to adapt the energetically expensive nervous system to environmental changes (Eriksson et al., 2019), optimizing the ratio between costs and benefits (Niven & Laughlin, 2008). We presently experience a rapid and overwhelming environmental change caused by human activities (Elhacham et al., 2020), some of them also creating artificial sources of geosmin. Especially the drinking water network (Bristow et al., 2019), agriculture (Lu et al., 2003) and aquaculture (Davidson et al., 2020) have the potential to release geosmin into the environment. Additionally, the anthropogenic climate change can enhance cyanobacterial blooms in natural water bodies (Paerl et al., 2016) as another source of geosmin (Chung et al., 2016). Such environmental changes can decrease the informational value of an odorant for an organism and can therefore be detrimental for its reproduction or survival. Experience-dependent plasticity in the olfactory system of insects is one way to counteract such detrimental effects by allowing quick modulations in an everchanging environment.

## MATERIAL & METHODS

### Fly stocks

Flies were reared in plastic vials on standard cornmeal medium and transferred to new vials every two weeks. Fly vials were kept in an incubator (Economic Premium, Snijders Scientific, Tilburg, Netherlands) under a 12h/12h light/dark cycle at 25°C and 70% humidity. Fly lines used for crossings or experiments are shown in the key resource table. Detailed genotypes used for each experiment are shown in the supplementary information (Table S2).

### Odor exposure

Flies were periodically exposed to either the odor or to mineral oil in a dedicated odor exposure setup. The setup consisted of four threaded glass bottles, which were connected with each other and with the air supply system in the lab via flexible Teflon tubes. The airstream entered the setup through a flow meter (Brooks Instrument, Dresden, Germany), which adjusted the air flow to one liter per minute. The first bottle (250 ml, Schott, Jena, Germany) contained water and an air diffusor to humidify the airstream. An electronically controlled valve (3/2-way mini electric solenoid valve M5, 24V, Pneumatikatlas, Staufenberg, Germany), which switched periodically between two different directions was installed behind the water bottle. One direction was connected with a smaller glass bottle (100 ml, Fischer Scientific, Schwerte, Germany) that contained 1 ml of odor solution (at a concentration of 10^−3^ diluted in mineral oil) or mineral oil. The other direction was connected to another 100 ml bottle containing 1 cm of standard cornmeal medium in the bottom and the flies. A fourth bottle (100 ml), which contained activated charcoal, was connected behind the fly bottle to bind the odor molecules before the airstream left the setup through a second flow meter that checked the whole system for tightness. A control unit for the valve was designed by J. Wilde and D. Veit (Jena). It used a program written in the programming language Python (by J. Cramer and D. Veit, Jena) that automatically switched between the two directions. Several setups could be connected to the control unit and operated simultaneously. Freshly emerged flies in a mixed population that contained males and females were transferred into the fly bottle. Flies were supplied with fresh air only during the first day to get used to the environment. The exposure started on the second day. During the exposure, flies periodically received the odor/mineral oil for 1 min, followed by 5 min of fresh air. The exposure lasted for four days and afterwards the flies were used for experiments (Fig. 1B). The setup contained either 6-20 flies (equal ratio of females and males) or 130-150 flies (100-120 females, 30 males), which were used for morphological/physiological or behavioral experiments, respectively. The exposure setups were placed in a breeding chamber (25°C, 70% humidity and a 12 h /12 h light/dark cycle).

### 2-photon photoactivation

For in vivo photoactivation experiments, 5-day-old flies that express photoactivatable GFP (C3PA-GFP) in subsets of projection neurons, local interneurons or glia cells were used. The flies were previously exposed to either the odor or to mineral oil and subsequently dissected largely following the description in Strutz et al. (Strutz et al., 2012). For the dissection, we pushed the cervical region of an anesthetized fly into a slit of a copper plate (Athene slot diaphragm, 125 μm slot, Plano, Wetzlar, Germany) that was glued to a custom made Plexiglas mounting stage. After this procedure, only the head was on top of the copper plate, while the thorax and abdomen were below it. We immobilized the head by pressing a fine needle (minutiens 0.10 mm, Austerlitz Insect Pins, Slavkov u Brna, Czech Republic) on top of the proboscis. The fine needle was then fixed with beeswax. To further stabilize the head, we glued its backside to the copper plate using 3-component dentist glue (Protemp™ II, 3M, Neuss, Germany). A fine tetrode wire (Redi Ohm 800, H.P. Reid Inc., Palm Coast, USA) that was fixed to a plastic plate with beeswax at both ends was inserted into the ptilinal suture. The plastic plate was slowly pushed forward by screws in the Plexiglas mounting stage to displace the antennae slightly to the front. A plastic plate with a hole was glued to a piece of polyethylene foil. A smaller hole was punched into the foil and the plastic plate was then placed on top of the fly’s head, so that the hole in the foil revealed the head. The space between the edges of the foil and the head capsule of the fly was sealed with 2-component silicon (Kwik-Sil™, World Precision Instruments, Sarasota, USA). A droplet of Ringer’s solution (NaCl: 130 mM, KCl: 5 mM, MgCl2: 2 mM, CaCl2: 2 mM, Sucrose: 36 mM, HEPES-NaOH (pH 7.3): 5 mM) was added on top of the head. Subsequently, the head capsule was opened dorsally with a fine scalpel (Micro Knive, Fine Science Tools, Heidelberg, Germany). Fat, glands and tracheae were removed with fine forceps to allow optical access to the brain. The mounting stage with the fly was then placed on the stage of the 2-photon microscope.

For the acquisition of PN scans, brains of dissected flies that express photoactivatable GFP in a subset of PNs (GH146-Gal4;UAS-C3PA-GFP) were scanned with a ZEISS LSM 710 NLO confocal microscope (Carl Zeiss, Jena, Germany) equipped with an infrared Chameleon Ultra diode-pumped laser (Coherent, Santa Clara, USA). The microscope was controlled by the ZEN microscopy software (Carl Zeiss, Jena, Germany) and operated in the non-descanned mode (NDD). For the photoactivation procedure, a 40x water immersion objective (W Plan-Apochromat 40x/1.0 DIC VIS-IR, Carl Zeiss, Jena, Germany) was used. A first stacked scan of the AL was obtained with a laser wavelength of 925 nm and a laser power of 30-40% to identify the target glomerulus. If necessary, the laser power was increased throughout the stack to compensate for decreased fluorescent signals in lower tissue layers. A precise region of interest was then placed on a layer in the middle of the glomerulus and continuously illuminated for 15 min at a wavelength of 760 nm and at 3-5% laser power. This was done for either one brain hemisphere or both. Afterwards, flies were kept in a dark humidified chamber for 25 min to allow photoconverted GFP molecules to diffuse within the PNs. Subsequently, the fly was sacrificed to reduce movements. A final scan was carried out with a slice interval of 1 μm, a pixel resolution of 1024 × 1024 and at the initial wavelength of 925 nm but with much less laser power (6-20%) to visualize only those PNs that contain photoconverted GFP molecules. The photoactivation approaches for glia cells and LNs were very similar to the PN procedure. To label glia cells, we used flies that express photoactivatable GFP in glia cells and mtdTomato in a subset of PNs (UAS-C3PA-GFP;Repo-Gal4/GH146-QF,QUAS-mtdTomato) and scanned their brains with a 63x water immersion objective (W Plan-Apochromat 63x/1.0 VIS-IR, Carl Zeiss, Jena, Germany), extended the photoactivation time to 25-30 min and created final scans with a 0.77 μm interval and 624 × 624 pixel resolution. The mtdTomato signal was recorded in a separate channel and allowed us to find the target glomerulus. For the photoactivation of LNs, we used flies that express photoactivatable GFP in different subsets of LNs and mtdTomato in a subset of PNs (“LN-Gal4” + UAS-C3PA-GFP; GH146-QF, QUAS-mtdTomato). Instead of targeting the area of the glomerulus, we put a small region of interest on a single LN soma to photoconvert GFP molecules in individual LNs. For each “LN-Gal4” driver line, we always tried to pick LN somata in the same region of the cell body cluster to minimize the chance of photoactivating LNs of completely different subtypes. To ensure that photoconverted GFP molecules diffuse throughout the entire LN, we extended the photoactivation time to 25-30 min and the time in the humidified chamber to at least 25 min. Final scans were done with the 63x objective, a slice interval of 0.77 μm and a pixel resolution of 624 × 624. The mtdTomato signal was used for orientation.

### 3D reconstructions

For 3D reconstructions, the acquired scans were processed with AMIRA 5.6 software (Fei Visualization Sciences Group) using the LabelField module and the semi-automated reconstruction plugin hxskeletonize (Evers et al., 2005; Schmitt et al., 2004) to reconstruct glomeruli and neurons, respectively. Glomeruli and somata were labeled manually on every second slice of the stack and later interpolated. The interpolation was checked for precision and if necessary corrected manually. The volumetric data of the assigned labels were measured with the MaterialStatistics module. Neurons were traced and their diameter was adjusted manually with the GraphEditor tools of the hxskeletonize module. Complete neurons were only reconstructed for demonstration purposes. For PN data, we reconstructed only the axonal branches beginning at the point at which PNs enter the LH. For glomeruli, which are innervated by more than one PN, we could only create one common reconstruction for all PNs that were labeled by the photoactivation procedure as we could not discriminate between individual PNs. Only LN parts that innervate the target glomeruli were reconstructed for the acquisition of LN data. Length and volume measurements were extracted from the properties panel of the DisplaySkeletonGraph module. The numbers of axonal terminals and boutons were counted manually.

### Volume quantifications of cell processes in glomeruli

To assess the volume of PN dendrites inside the DA2 glomerulus, we used stacks from our photoactivation experiment. First, we labeled the glomerulus with the LabelField module in AMIRA 5.6 and assigned this selection to our first material. We assigned the area outside of the glomerulus to a second material. Then we locked the second material and used the threshold tool to select voxels of certain brightness across the whole image stack. For PNs, we used a threshold of 70-255, which excludes voxels with a grey value less than 70. This range selected voxels that covered the signal of our datasets well. By assigning the selection to a third material, the selection got subtracted from the glomerulus volume while parts of the PNs outside of the glomerulus were untouched because they were locked. Consequently, we obtained three materials two of which contained the information about the total glomerular volume and the volume of the PN dendrites inside this particular glomerulus. The volumetric values were extracted with the MaterialStatistics module. Scans with PN dendrites occupying less than 25% or more than 75% of the total glomerulus volume were considered as underexposed or overexposed, respectively, and were excluded from the analysis. We had to exclude some of the scans as they were not initially planned to resolve the PN dendrites in the best possible way, but to yield sufficient resolution in the region of the LH to conduct 3D reconstructions and for precise counting of the PN somata. The same approach was used to assess OSN, LN and glia cell data, except that we did not have to exclude datasets as we exposed properly for the region of the AL. For the acquisition of stacks for our OSN experiment, we used flies that express neoGFP under the control of Orco-Gal4 (UAS-neoGFP; Orco-Gal4) and scanned their brains with the setup described in the 2-photon photoactivation section. To select OSN axons, we used a threshold of 800-4095. The procedure we used to acquire stacks for our LN and glia cell experiment is described in the 2-photon photoactivation section. The threshold for glia selection was 70-255. The threshold for LN selection differed slightly between the different LN lines because of variations in the expression levels (GH298: 20-255; NP3056: 30-255; NP2426, HB4-93: 40-255).

### Oviposition assay

A schematic illustration of the oviposition assay is showninfigure 6D.Weusedglasspetridishes(13.5 cm wide, 2 cm high) as containers and smaller plastic petri dishes (3.3 cm wide, 1 cm high) as choice plates. Standard cornmeal medium was poured inside the small plastic petri dishes up to a height of 0.5 cm. After cooling down and solidifying, we used a 200 ml pipette tip to punch holes in the center of the food surface. Either 10 μl of odor (geosmin 10^−3^) or mineral oil were pipetted into the hole, which was then covered by a circular piece of filter paper. The choice plates were then placed with a distance of 2 cm from each other in the center of the glass petri dishes and arranged randomly in different setups. Ten female flies, which were previously in the exposure setup together with males to have the opportunity to mate, were added to the setup, which was then closed with the glass lid. We started the experiments in the afternoon and terminated them after 24 h. For the duration of the experiments, the setups were placed in a breeding chamber (25°C, 70% humidity and a 12 h /12 h light/dark cycle). After 24 h, the entire setups were put into a freezer to anesthetize the flies. Afterwards, the number of eggs on each choice plate was counted. Oviposition indices (OI) were calculated (OI = (O-C)/T), with O as the amount of eggs on the odor plate, C the amount of eggs on the control plate and T the total amount of eggs on both plates. Plastic petri dishes were discarded and glass petri dishes were cleaned for further use after the experiment. Dependent on how many flies survived the exposure period, 7-11 setups were run simultaneously in one trial. The presented oviposition indices represent the pooled data of three trials of three subsequent weeks (1 trial per week).

### T-maze assay

We used plastic t-tubes, which were 8 cm wide and 5.2 cm high, as T-maze assays. We plugged Eppendorf reaction tubes (1.5 ml, Greiner Bio-One GmbH, Frickenhausen, Germany), which were cut at their tip, into both ends of the horizontal part of the t-tube. A round piece of filter paper was placed into each Eppendorf tube. 3 μl of either odor (geosmin 10^−3^) or mineral oil were pipetted onto the filter paper. In one set of experiments, we added a cube of approximately 0.125 μm^-3^ standard cornmeal medium in both Eppendorf tubes right on top of the filter paper. The Eppendorf tubes were closed with their snap-on lid just before we added the flies to the setup. 10 female flies, which were previously in the exposure setup together with males, were shortly anesthetized in a freezer at -20°C and then put into the setup via the third opening that was not plugged yet with an Eppendorf tube. A funnel was used for this procedure. After flies were added, we closed the third opening with a plastic plug. Setups were placed in a breeding chamber (25°C, 70% humidity and a 12 h /12 h light/dark cycle) in standing position (plug on the bottom, as in Fig. 6A). In our arrangement, two neighboring setups had the odor and mineral oil on opposite sides. After 40 minutes, we terminated the experiment and counted the number of flies in each side of the two choice arms. Preference indices (PI) were calculated (PI = (O-C)/T), with O as the amount of flies on the odor side, C the amount of flies on the control side and T the total amount of flies in the setup (T=10). Dependent on how many flies survived the exposure period, 7-11 setups were run simultaneously in one trial. The presented preference indices represent the data of three trials of three subsequent weeks (1 trial per week). Previous investigations have shown that the morphological effects caused by long-term exposure can be completely reversed within 2 days (Chodankar et al., 2020; Kidd & Lieber, 2016; Kidd et al., 2015; Sachse et al., 2007). For this reason, we wanted to test our flies as soon as possible after they left the exposure setup and did not starve them for our T-maze assays. However, satiated flies are unmotivated and often fail to enter funnel traps that are usually used in T-maze assays. Consequently, we only cut the Eppendorf tubes at their tip and did not build a funnel there (easier access to the odor source). Additionally, we counted a choice already when a fly left the center part of the setup, without the need of actually entering one of the two Eppendorf tubes (see Fig. 6A).

### Immunostaining protocol

For immunostaining experiments, we used 5-day-old female flies expressing GFP tagged to Bruch-pilot-Short (Brp-Short) in subsets of local interneurons. The flies were previously exposed to either the odor or to mineral oil and subsequently dissected. For the dissection, the abdomen of an anesthetized fly was pinned to the center of a silicon-covered (Sylgard® 184, Dow Corning Corporation, Midland, USA) glass petri dish with a minutien pin. The entire fly was then covered with a drop of Ringer’s solution (NaCl: 130 mM, KCl: 5 mM, MgCl2: 2 mM, CaCl2: 2 mM, Sucrose: 36 mM, HEPES-NaOH (pH 7.3): 5 mM). The proboscis was cut off with fine micro scissors at its base, thus creating an opening in the head. Afterwards, the margin of the hole was grabbed with fine forceps on both sides and pulled apart. In some cases, this procedure already exposed the entire brain, which was still in its natural position, but covered by tracheae and fat body tissue. Subsequently, remaining parts of the cuticle, tracheae and fat body lobes were removed from the brain. After this, the cervical connective was severed and the cleaned brain was then transferred with fine forceps in a 500 μl Eppendorf reaction tube containing 500 μl of Ringer’s solution. Up to 18 brains were collected in one tube. For fixation of brains, the Ringer’s solution in the tube was replaced with phosphate buffer saline (PBS, 1x) containing 4% paraformaldehyde (PFA) and 0.1% Triton™ X-100 (Sigma-Aldrich, Taufkirchen, Germany). The tube was taped horizontally on an aluminum cooling rack in a plastic box containing ice, which was then placed on a shaker for 30 min. Afterwards, the brains were washed (3x shortly, 3x for 15 min at room temperature on a shaker) in PBS containing 0.1% Triton™ X-100 (PBST). The PBST was then replaced with blocking solution (PBST + 2% normal goat serum) and the tube was kept on the shaker at room temperature for another 30 min. Subsequently, the blocking solution was replaced with blocking solution containing the diluted primary antibodies. The brains were then incubated for approximately 48 h on a shaker at 4°C. The incubation was followed by the same washing procedure as before (PBST, 3x quickly, 3x for 15 min at room temperature on a shaker). The PBST was then replaced by blocking solution containing the secondary antibodies followed by incubation for another 48 h on a shaker at 4°C. After the incubation with the secondary antibodies, the brains were washed in PBST as mentioned before.

For mounting, we cut a cover slip in half (using a glass cutter) and glued each of these halves with nail polish on a microscope slide with a small gap between them. The brains were then transferred inside the gap with a glass Pasteur pipette. Fine forceps were used to arrange them properly. Once the brains were in position, the remaining PBST was soaked up with paper tissue and the brains were covered with VectaShield (Vector Laboratories, Burlingame, USA). A second cover slip was placed above the gap and sealed all around with nail polish. Microscope slides were stored in a fridge in darkness and at 4°C. We used rabbit anti-GFP (1:500, Invitrogen, Thermo Fisher Scientific) and mouse anti-bruchpilot (nc82, 1:30, Developmental Studies Hybridoma Bank) as primary antibodies and Alexa Fluor 488 goat anti-rabbit (1:250, Invitrogen, Thermo Fisher Scientific) and Alexa Fluor 633 goat anti-mouse (1:250, Invitrogen, Thermo Fisher Scientific) as secondary antibodies.

### Semi-automatic quantification of Brp-Short puncta

To assess the number of Brp-Short puncta of LN subpopulations within glomeruli, we used flies that express GFP coupled to Brp-Short in LNs (GH298-Gal4, NP2426-Gal4). Flies were previously exposed to either the odor or to mineral oil and subsequently dissected. The dissection and immunostaining (against nc82 and GFP) were conducted as mentioned above. We used the online Nyquist calculator from SVI (https://svi.nl/NyquistCalculator) to calculate the Nyquist rate for our particular microscopy setup and experiment. Since the calculation is dependent on the fluorescence properties of the used fluorophores, we calculated the Nyquist rate accordingly to the dye we used for staining Brp-Short (and not nc82) as we wanted to get the highest possible resolution of these structures. The preparations were then scanned with a Zeiss LSM 880 confocal microscope in two separate channels with at least the calculated resolution (0.05 × 0.05 × 0.16 μm). We obtained oversampled datasets, which allowed us to enhance the signals in our datasets by performing deconvolutions later on.

The acquired scans were imported into AMIRA 5.6 software (Fei Visualization Sciences Group).The signal of the nc82 staining was used to find the glomeruli of interest (DA2 and DA4l). We then manually labeled each of the two glomeruli on a separate LabelField using the LabelField module. Next, we used the Arithmetic tool with the expression A*(B>0), where A is the Brp-Short channel of our scan and B is the LabelField. Using this expression turns everything that was not labeled to black and only keeps the Brp-Short signal inside the labeled glomerulus. The resulting image stack was then cropped maximally to increase the speed of the subsequent deconvolution. This procedure was used for both glomeruli so that we obtained two small image stacks for each of our initial scans containing only the Brp-Short signal of two single glomeruli. Stacks were saved as TIFF image sequences.

Next, we imported these cropped stacks into ImageJ (Rueden et al., 2017; Schindelin et al., 2012).We used the Diffraction PSF 3D plugin to create a hypothetical point spread function (PSF) for our microscopy setup and experiment. A separate PSF was calculated for each image stack to fit the voxel dimensions of the crop. Next, we used the Iterative Deconvolve 3D plugin with suggested default settings. The deconvoluted image stacks were saved as TIFF image sequences for documentation and subsequently processed with the Interactive H_Watershed plugin of the SCF plugin collection. Watershedding splits clusters of densely packed Brp-Short puncta into smaller pieces, which otherwise would be counted as one puncta by conventional thresholding procedures. We randomly selected a few reference image stacks and interactively evaluated which parameters fit our data best. We then used these parameters consistently for all of our datasets (seed dynamics: 10% of maximum value, intensity threshold: 5% of maximum value, peak flooding: 80%, splitting al-lowed). The region mask of the watershed was exported to create a binary image stack in which Brp-Short puncta are pure white and everything else is pure black. The resulting stacks were then analyzed using the 3D Object Counter v2.0 plugin (Bolte & Cordelieres, 2006). Since we analyzed binary image stacks, we could neglect the threshold slider of the plugin. To select values for the size filter, we calculated an approximation of the volume of one Brp-Short punctum. With a Brp-Short punctum diameter of ∼0.6μm (Mosca & Luo, 2014) and the assumption that these structures are spherical, we hypothesized that one Brp-Short punctum has a volume of ∼0.113 μm^-3^. We picked a voxel count, which corresponds to 1/4 of the volume of a Brp-Short punctum for the minimum size filter, to exclude too small structures from the quantification. The maximum size filter was left at its default settings. The results in the statistics table were then copied into an Excel sheet for further analyzes.

### Semi-automatic quantification of mitochondria

We used flies that express GFP in mitochondria of OSNs, which innervate the DA2 glomerulus (Or56a-Gal4, UAS-mitoGFP). Flies were previously exposed to either the odor or to mineral oil and subsequently dissected as described for the 2-photon photoactivation experiments. They were sacrificed immediately after dissection and their esophagus was removed to minimize movements during the scan. The Nyquist rate was calculated as described above and oversampled scans (voxel size: 0.051μm x 0.051μm x 0.271μm) of the ALs were conducted with the microscopy setup that we described for the 2-photon photoactivation experiments. The signal of mitochondria alone was sufficient to see glomerular borders clearly. Scans were processed as described for the quantification of Brp-Short puncta. We changed the value for seed dynamics in the Interactive Watershed plugin to 15% of the maximum value because this way the results reflected the signal in the scans more precisely. We set the minimum size filter of the 3D Object Counter plugin to 100 voxel as this value yielded results that matched manual counting of mitochondria.

### 2-photon calcium imaging

To quantify neuronal signaling of OSNs and PNs in the DA2 glomerulus, we used flies that express the calcium sensor GCaMP in these neurons (OSNs: Orco-Gal4, UAS-syp-GCaMP3; PNs: GH146-Gal4, UAS-GCaMP6f). Prior to the imaging experiment, flies were exposed to either geosmin or to min-eral oil and subsequently dissected as described for the 2-photon photoactivation experiments. An electronically controlled odor delivery system was utilized to guide airborne odor molecules to the antenna of dissected flies (as described in Mohamed et al. (2019). In brief, the delivery system consists of flexible Teflon tubes that guide two converging air-streams (0.5 l/min each) to the antenna. A solenoid valve, which is controlled by the LabVIEW software (National Instruments, Austin, USA), is installed in one of these airstreams and switches between one tube transporting pure air and one tube that enters a 50 ml glass bottle (Schott, Jena, Germany) containing 1 ml of diluted odorant (geosmin: 10^−3^-10^−7^; all other odorants: 10^−3^). We used the same microscopy setup that we described for the 2-photon photoactivation experiments to acquire time series. The fluorophore of GCaMP was excited by a laser wavelength of 925 nm. An argon laser with a wavelength of 488 nm was only used for the PN responses in the LH, due to technical problems with the 2-photon laser prior to the experiment. For each fly, we conducted the experiment on one side of the AL. Prior to acquiring the time series, we obtained a z-stack to get an overview of the AL and to find our target glomeruli. We selected two focal planes that covered the D, DA2, DA4l, DM1 and DM2 glomeruli. In both focal planes, we exposed the dissected fly to pure air (blank), mineral oil, 1-hexanol, 3-hexanone, benzaldehyde, ethyl acetate and geosmin. The order of the different odorants was randomized for each fly. However, the different concentrations of geosmin were always presented in sequence and in an ascending order. Each time series had a length of 10 s and consisted of 40 frames (framerate of 4 Hz). The odor was presented after 2 s and continued for 2 s. We paused after each time series to flush the tubes and to wait for the fluorescence to get back to baseline. The pixel dimension of a single frame was 248 × 250. Time series were aligned in ImageJ to compensate for movements by using the StackReg plugin (Thévenaz et al., 1998). The ROI Manager was used to label glomeruli of interest and to extract the average brightness value of pixels within the selected areas across the entire time series. These values were imported into RStudio (RStudio Team, 2016) to calculate ΔF/F [%]. For this, the mean values of the first 8 frames (= base fluorescence) were first subtracted from all values of the time series to calculate ΔF. ΔF was then divided by the base fluorescence to calculate ΔF/F and subsequently multiplied by 100 to obtain ΔF/F [%]. The mean of ΔF/F [%] was calculated by averaging those 13 frames of the time series that covered the maximum response. Frames 9-21, 13-25 or 17-29 were used. This time window was kept constant within an experiment (e.g. all values of the mean ΔF/F [%] of Orco-Gal4, UAS-syp-GCaMP3 responses result from frames 17-29). The exact time windows used for each experiment are indicated as black bars in Fig. S3.

### Figures and statistical analyses

Graphs were made with the programming language R (R Core Team, 2018) and RStudio (RStudio Team, 2016) using the packages ‘ggplot2’ (Wickham, 2016) and ‘reshape’ (Wickham, 2007) and exported to the Adobe Illustrator CS5 software, which we used to compile figure plates. Individual data points are shown in graphs wherever the space permitted it. Boxplots show the median (horizontal line), the 25th and 75th percentile (lower and upper border of the box, respectively) and the lowest and highest values that lie within 1.5 times of the inter-quartile range of the box (whiskers). Error bars in the dose response curves indicate the standard deviation of the mean. Representative maximum intensity projections were made and single planes of z-stacks were extracted with ImageJ and cropped and adjusted for brightness and contrast in the Adobe Photoshop CS5 software. For statistical analyses, we used Microsoft Excel 2010 (one-sample student’s t test, two-sample student’s t test) and IBM SPSS Statistics 25 (two-way repeated measures ANOVA with Tukey post-hoc test). The parametric two-sample ttest was chosen to evaluate whether the mean of two groups differs. The reasons for this choice were the large sample size in most experiments, which makes the parametric t-test robust even in cases where the data were not normally distributed, and the symmetric distribution of the data points. The statistical tests used for the data in each figure are mentioned in the respective figure legends. Outliers were not excluded from statistical analyses. The following significance levels were used throughout this study: * p < 0.05, ** p < 0.005, *** p < 0.0005.

## ACKNOWLEDGEMENTS

We would like to thank Silke Trautheim for her excellent support for fly rearing and technical assistance. We thank members of the Sachse and Hansson labs for discussions and comments on the study. This study was funded by the Max Planck Society and was part of the International Max Planck Research School (IMPRS) ‘Chemical Communication in Ecological Systems’.

## AUTHOR CONTRIBUTION

Conceptualization, S.S.; Methodology, B.F., V.G., and S.S.; Formal Analysis, B.F.; Investigation, B.F.; Writing – Original Draft, B.F.; Writing – Review & Editing, B.F., V.G., R.G.B., B.S.H., and S.S.; Visualization, B.F.; Supervision, S.S.; Project Administration, S.S.; Funding Acquisition, B.S.H., R.G.B. and S.S.

## DECLARATION OF INTEREST

The authors declare no competing interests.

## Notes

### Competing Interest Statement

The authors have declared no competing interest.

### Summary of Updates

Supplementary table S1 in the supplemental file was exchanged as the previous table was an old version.

