## Supplementary Information for "Experience-dependent plasticity of a highly specific olfactory circuit in *Drosophila melanogaster*"

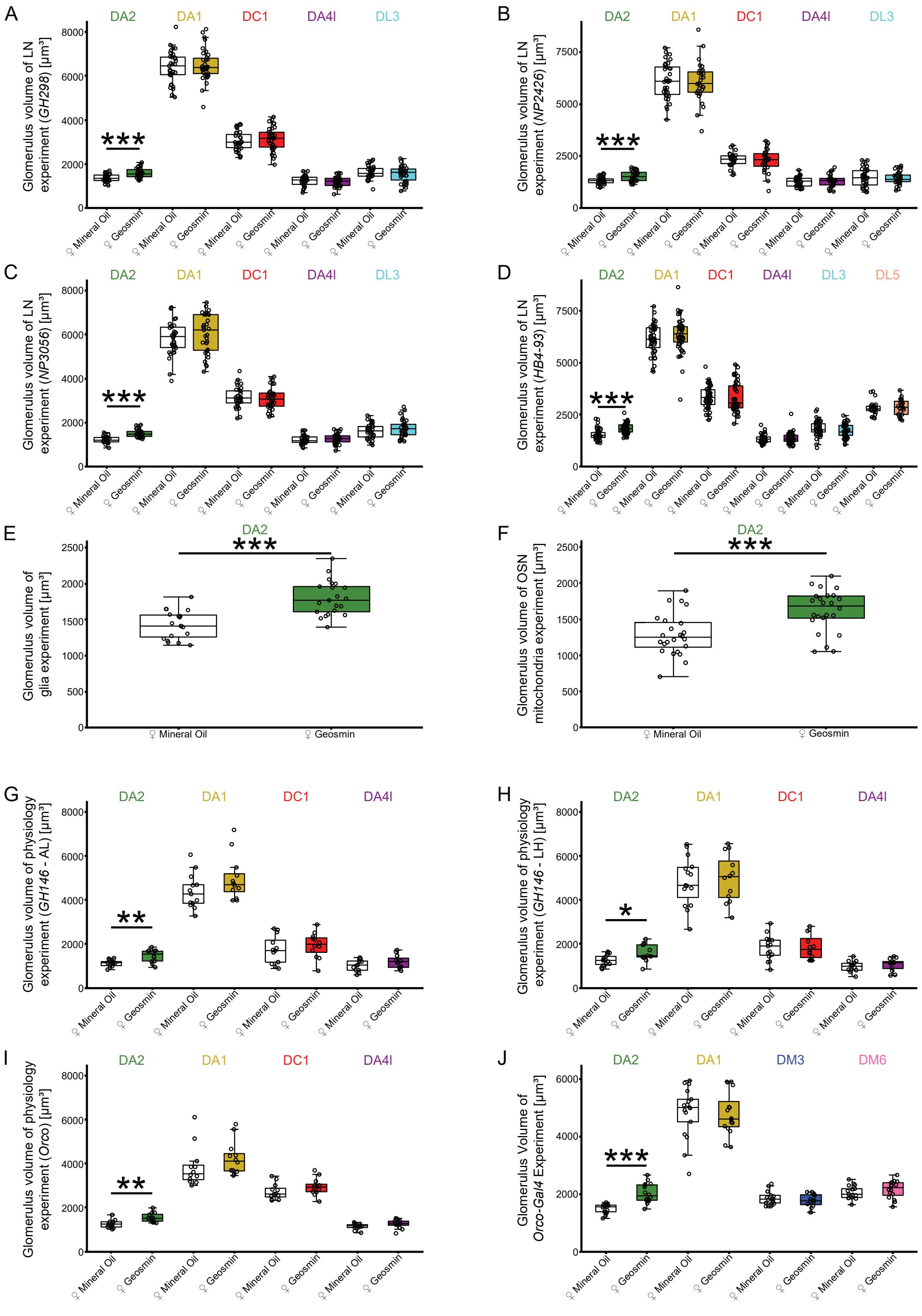


Fig. S1: Glomerular volume in different experiments. The DA2 glomerulus increased in size due to geosmin exposure, while the other measured glomeruli were unaffected. A, GH298- Gal4 experiment (n = 31-34). B, NP2426-Gal4 experiment (n = 26-34). C, NP3056-Gal4 experiment (n = 30-35). D, HB4-93-Gal4 experiment (DL5: n = 22-23; DA2, DA1, DC1, DA4l, DL3: n = 45-47). E, Glia experiment (n = 19-21). F, OSN mitochondria experiment (n = 24). G, GH146- Gal4 physiology experiment (AL) (n = 11-13). H, GH146-Gal4 physiology experiment (LH) (n = 12-16). I, Orco-Gal4 syp-GCaMP physiology experiment (n = 12). J, Orco-Gal4 experiment (n = 16-17). Statistical test: two-sample student’s t test. Significance levels: * $p < 0.05$, ** $p < 0.005$, *** $p < 0.0005$.

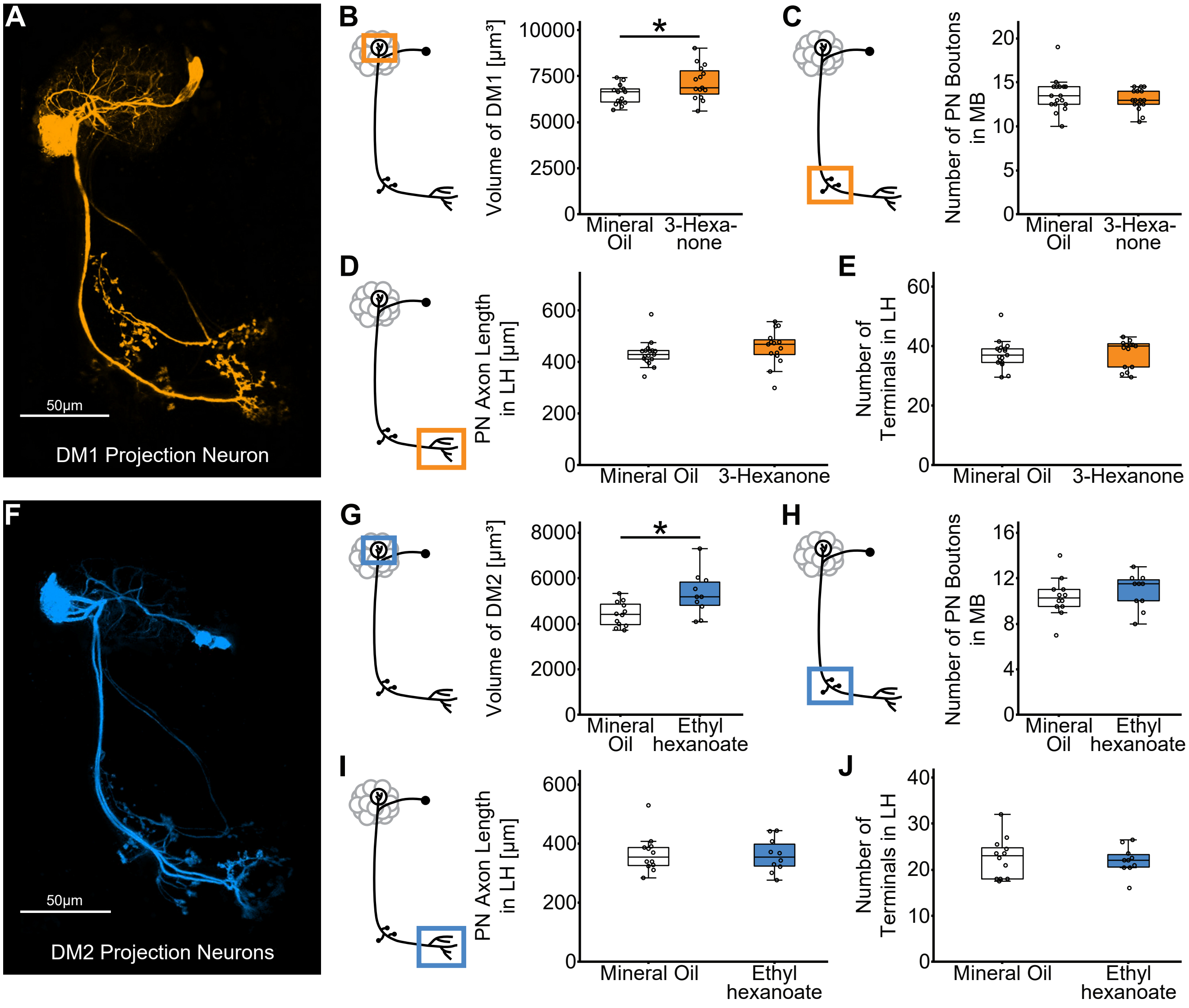


Fig. S2: Long-term exposure of *D. melanogaster* females to attractive odors leads to a volumetric increase of their associated glomerulus. A, representative z-stack of the sole projection neuron innervating the DM1 glomerulus. B, exposure to 3-hexanone leads to a volumetric increase of the DM1 glomerulus in females (n = 15). C, the number of projection neuron boutons in the mushroom body is unaltered after 3-hexanone exposure (n = 16-17). D, E, the axonal length and axonal terminals of projection neurons in the lateral horn are not affected by the 3-hexanone exposure (n = 15-17). F, representative z-stack of the two projection neurons innervating the DM2 glomerulus. G, exposure to ethyl hexanoate leads to a volumetric increase of the DM2 glomerulus in females (n = 10-12). H, the number of projection neuron boutons in the mushroom body is unaltered after ethyl hexanoate exposure (n = 10-12). I, J, the axonal length and axonal terminals of projection neurons in the lateral horn are not affected by the ethyl hexanoate exposure (n = 10-12). Statistical test: two-sample student's t test. Significance levels: * $p < 0.05$. Abbreviations: LH, lateral horn; MB, mushroom body, PN, projection neuron.

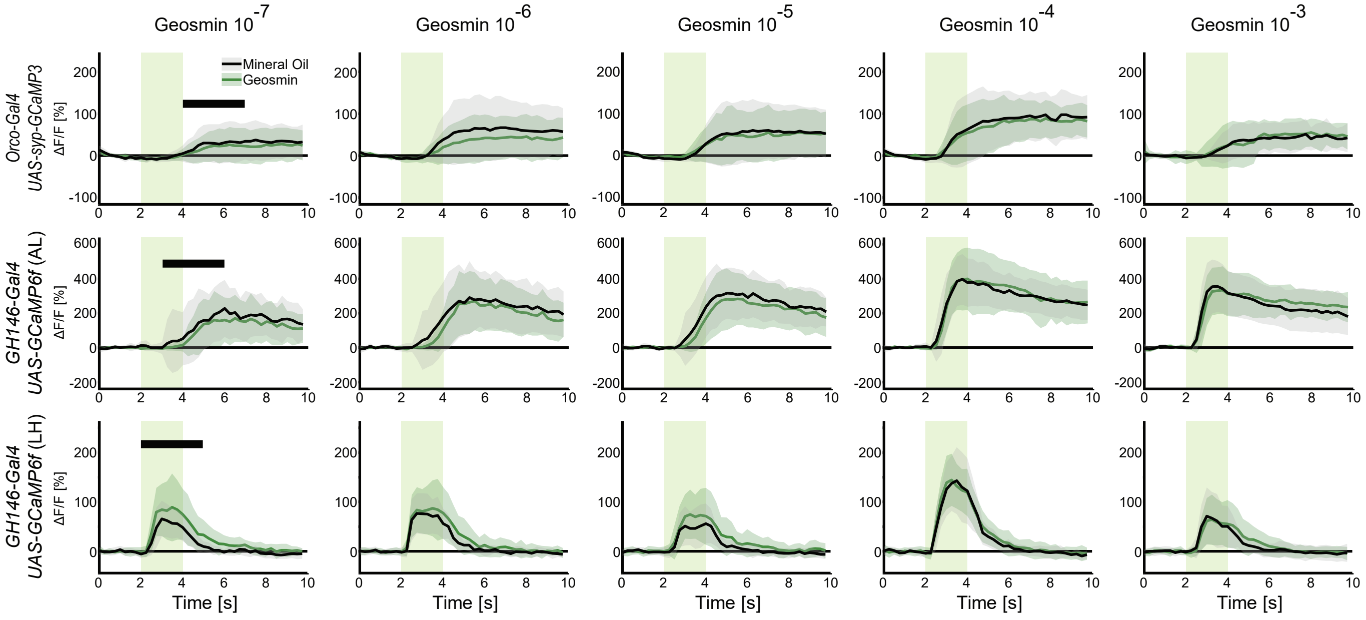


Fig. S3: Average time traces of calcium imaging responses of different DA2 neurons to different geosmin concentrations. Black lines show the average responses of mineral oil exposed flies and green lines show the average responses of geosmin exposed flies. The standard deviation is depicted as a transparent shadow in the background of these lines (green, geosmin exposed; grey, mineral oil exposed). The green rectangle between 2-4 s symbolizes the odor puff. Black bars indicate the 3 s time window that was used to calculate the average maximum response in each animal to acquire the data shown in Fig. 7. We chose the time window in a way that it is still close to the odor puff but also covers the highest responses. This time window was kept constant for a given experiment (e.g. for the GH146-Gal4, UAS-GCaMP6f (AL) experiment we always used 3-6 s). Sample sizes: Orco-Gal4, UAS-syp-GCaMP3, n = 12; GH146-Gal4, UAS-GCaMP6f (AL), n = 12-13; GH146-Gal4, UAS-GCaMP6f (LH), n = 17.

Table S1: Fly genotypes that were used for each figure.

| Figure | | Genotype |
|---|---|---|
| Fig. 1 | A-I | *yw; GH146-Gal4/CyO; UAS-C3PA-GFP/TM6B* |
| Fig. 2 | A-A1 | *yw/w1118; UAS-C3PA-GFP/BL; GH298-Gal4/GH146-QF,QUAS-mtdTomato* |
| | B-B1 | *NP2426-Gal4/w1118; UAS-C3PA-GFP/+; GH146-QF,QUAS-mtdTomato/(TM2/TM6B)* |
| | C-C1 | *w*/w1118; UAS-C3PA-GFP/BL; NP3056-Gal4/GH146-QF,QUAS-mtdTomato* |
| | D-D1 | *w+/w1118; UAS-C3PA-GFP/+; HB4-93/GH146-QF,QUAS-mtdTomato* |
| Fig. 3 | A, C-C5 | *NP2426-Gal4/w1118; UAS-C3PA-GFP/+; GH146-QF,QUAS-mtdTomato/(TM2/TM6B)* |
| | B-B5 | *yw/w1118; UAS-C3PA-GFP/BL; GH298-Gal4/GH146-QF,QUAS-mtdTomato* |
| | D-D5 | *w*/w1118; UAS-C3PA-GFP/BL; NP3056-Gal4/GH146-QF,QUAS-mtdTomato* |
| | E-E5 | *w+/w1118; UAS-C3PA-GFP/+; HB4-93/GH146-QF,QUAS-mtdTomato* |
| Fig. 4 | A-E | *NP2426-Gal4/+ ; +/+; UAS-BRP-short-GFP/(TM2/TM6B)* |
| | F-I | *+/yw; +/(CyO/BL); GH298-Gal4/UAS-shortBRP-GFP* |
| Fig. 5 | A-B | *+; UAS-neoGFP; Orco-Gal4* |
| | C-F | *+; CyO/+; Or56a-Gal4,UAS-MitoGFP/TM6B* |
| | G-H | *+/ w1118; UAS-C3PA-GFP/BL; repo-Gal4/GH146-QF,QUAS-mtdTomato* |
| Fig. 6 | A-E | Wild-type Canton S |
| Fig. 7 | A | *yw/+; UAS-syp-GCaMP3/CyO; Orco-Gal4* |
| | B-C | *yw/w1118; GH146-Gal4/CyO; UAS-GCaMP6f* |
| Fig. S1 | A | *yw/w1118; UAS-C3PA-GFP/BL; GH298-Gal4/GH146-QF,QUAS-mtdTomato* |
| | B | *NP2426-Gal4/w1118; UAS-C3PA-GFP/+; GH146-QF,QUAS-mtdTomato/(TM2/TM6B)* |
| | C | *w*/w1118; UAS-C3PA-GFP/BL; NP3056-Gal4/GH146-QF,QUAS-mtdTomato* |
| | D | *w+/w1118; UAS-C3PA-GFP/+; HB4-93/GH146-QF,QUAS-mtdTomato* |
| | E | *+/ w1118; UAS-C3PA-GFP/BL; repo-Gal4/GH146-QF,QUAS-mtdTomato* |
| | F | *+; CyO/+; Or56a-Gal4,UAS-MitoGFP/TM6B* |
| | G-H | *yw/ w1118; GH146-Gal4/CyO; UAS-GCaMP6f* |
| | I | *yw/+; UAS-syp-GCaMP3/CyO; Orco-Gal4* |
| | J | *+; UAS-neoGFP; Orco-Gal4* |
| Fig. S2 | A-J | *yw; GH146-Gal4/CyO; UAS-C3PA-GFP/TM6B* |
| Fig. S3 | | *yw/+; UAS-syp-GCaMP3/CyO; Orco-Gal4* |
| | | *yw/w1118; GH146-Gal4/CyO; UAS-GCaMP6f* |

Table S2: Glomerular volume increase of all experiments in which we measured the glomerular volume.

| Glomerulus | Experiment | Sample sizes | Average volume of DA2 [µm³] | | | | Average volume difference [µm³] | Growth [%] |
|---|---|---|---|---|---|---|---|---|
| | | | Geosmin exposed | ± SD | Mineral oil exposed | ± SD | | |
| DA2 | PN photoactivation | 23, 26 | 2718,07 | 577,05 | 2121,20 | 342,58 | 596,87 | 28,14 |
| | OSN innervation | 16, 17 | 2036,96 | 334,55 | 1504,57 | 159,86 | 532,39 | 35,39 |
| | GH298 LN photoactivation | 33, 33 | 1752,56 | 187,28 | 1417,57 | 247,44 | 334,98 | 23,63 |
| | HB4-93 LN photoactivation | 50, 53 | 1867,85 | 312,00 | 1598,03 | 277,09 | 269,82 | 16,88 |
| | NP2426 LN photoactivation | 29, 34 | 1657,34 | 264,84 | 1373,18 | 227,48 | 284,16 | 20,69 |
| | NP3056 LN photoactivation | 31, 34 | 1584,46 | 188,38 | 1318,05 | 198,78 | 266,41 | 20,21 |
| | Glia photoactivation | 19, 21 | 1797,91 | 241,30 | 1427,67 | 193,66 | 370,24 | 25,93 |
| | OSN mitochondria | 24, 24 | 1619,30 | 294,62 | 1287,63 | 289,07 | 331,68 | 25,76 |
| | GH146 physiology (AL) | 12, 13 | 1459,60 | 289,57 | 1123,35 | 149,82 | 336,26 | 29,93 |
| | GH146 physiology (LH) | 12, 16 | 1594,63 | 396,76 | 1257,57 | 249,85 | 337,06 | 26,80 |
| | OSN physiology | 12, 12 | 1549,05 | 216,09 | 1256,83 | 180,61 | 292,22 | 23,25 |
| DM1 | PN photoactivation | 15, 15 | 7151,58 | 922,72 | 6496,37 | 530,05 | 655,20 | 10,09 |
| DM2 | PN photoactivation | 10, 12 | 5283,34 | 1034,61 | 4418,54 | 536,38 | 864,80 | 19,57 |
